# The open bar is closed: restructuration of a native parasitoid community following successful control of an invasive pest

**DOI:** 10.1101/2019.12.20.884908

**Authors:** David Muru, Nicolas Borowiec, Marcel Thaon, Nicolas Ris, Madalina I. Viciriuc, Sylvie Warot, Elodie Vercken

## Abstract

The rise of the Asian chestnut gall wasp *Dryocosmus kuriphilus* in France has benefited the native community of parasitoids originally associated with oak gall wasps by becoming an additional trophic subsidy and therefore perturbing population dynamics of local parasitoids. However, the successful biological control of this pest has then led to significant decreases in its population densities. Here we investigate how the invasion of the Asian chestnut gall wasp *Dryocosmus kuriphilus* in France and its subsequent control by the exotic parasitoid *Torymus sinensis* has impacted the local community of native parasitoids.

We explored 5 years of native community dynamics within 26 locations during the rise and fall of the invasive pest. In an attempt to understand how mechanisms such as local extinction or competition come into play, we analyzed how the patterns of co-occurrence between the different native parasitoid species changed through time.

Our results demonstrate that native parasitoid communities experienced increased competition as the *D. kuriphilus* levels of infestation decreased. During the last year of the survey, two alternative patterns were observed depending on the sampled location: either native parasitoid communities were represented by an extremely limited number of species occurring at low densities, in some cases no native parasitoid species at all, or they were dominated by one main parasitoid: *Mesopolobus sericeus*. These two patterns seemed to correlate with the habitat type, *M. sericeus* being more abundant in semi-natural habitats compared to agricultural lands, the former known to be natural reservoirs for native parasitoids. These results highlight how the “boom-and-bust” dynamics of an invasive pest followed by successful biological control can deeply alter the structure of native communities of natural enemies.

This article has been peer-reviewed and recommended by *Peer Community in Zoology* https://doi.org/10.24072/pci.zool.100004

## Introduction

Biological invasions are defined as the introduction, establishment and expansion of populations outside of their native area. These two patterns seemed to correlate with the habitat type – *M. sericeus* being more abundant in semi-natural habitats compared to agricultural lands (see McMichael and Bouma 2000 for a review), the former known to be natural reservoirs for native parasitoids. More generally, invasions are known to have a diversity of direct and indirect negative effects on native ecosystems (see McGeoch et al. 2015 for a review of environmental impacts caused by invasion). In particular, they can deeply alter interspecific interactions and restructure native communities, often with negative consequences (Ricciardi and Isaac 2000, Ricciardi 2001, Carroll 2007). For instance, invasive predators, parasitoids or pathogens were proven to drastically reduce the size of resident prey or host populations (Daszak et al. 2000) or to compete with other species from the same trophic level (Hamilton et al. 1999, Grosholz 2002, see David et al. 2017 for a review of the impacts of invasive species on food webs).

While much research has focused on invasive top-consumers (predators, parasitoids, etc.), a more restrained amount of literature examines the impact of primary-consumer invasive species as a new resource for the native community (Carlsson et al. 2009). The situation where a successful invasive species becomes a resource for native species of a higher trophic level is referred to as a form of facilitation and the invasive species acts as a trophic subsidy (Rodriguez 2006). The ecological impact of invasive species acting as a trophic subsidy has been shown across a variety of model systems such as invasive macroalgae (Olabarria et al. 2009, Rossi et al. 2010, Suarez-Jimenez et al. 2017), some phytophagous insects (Barber et al. 2008, Girardoz et al. 2008, Jones et al. 2014, Haye et al. 2015, Herlihy et al. 2016, Noyes 2017), *Drosophila suzukii* (Mazzetto et al. 2016) or four invasive gall wasp species (Schonrogge and Crawley 2000). Hence, trophic subsidies, possibly in pair with changes in population densities of local populations (Eveleigh et al. 2007) may change the established dynamics within recipient communities through, for example, apparent competition (Settle and Wilson 1990, Holt and Bonsall 2017) or niche displacement (Mallon et al. 2007). Even when the invasion is only transient and therefore the trophic subsidy finally disappears, the consequences on native communities can be lasting on the recipient community. Indeed, recovery from the disappearance of the alien species is not systematic and may thus lead to further disequilibrium within the community (Courchamp et al. 2003). For example, Mallon et al. (2017) have recently reported a permanent niche displacement of native species caused by a failed invasion by *Escherichia coli* in soil microcosms, and referred to it as a “legacy effect”. Such lasting effects of transient invasions on species niche breadth and space for instance, may as well occur at the community level, thus impacting overall community structure by forcing species to modify the way they exploit available resources. However, empirical data on the response of native community dynamics to transient invasion remain scarce, as there is a lack of studies exploring multi-year community dynamics during the rise and fall of an invasive pest.

Classical biological control – i.e. the deliberate introduction of an exotic biological control agent to durably regulate a target (usually exotic) pest (Eilenberg et al. 2001) - can provide valuable empirical data on the dynamics of communities disturbed by two successive invaders, the pest and its introduced natural enemy. Indeed, during a biological control program, the dynamics of local communities are disturbed twice consecutively. Firstly, the arrival of the invasive pest decreases the level of primary resource and can alter the abundances of its native competitors and natural enemies (Jones et al. 2014, Haye et al. 2015, Herlihy et al. 2016). Then, the establishment of the biological control agent will again modify the community structure by strongly reducing the abundance of the invasive pest and potentially interacting with native species. Long-term direct and indirect effects of either the pest or its natural enemy on the recipient community have been documented (Henneman and Memmott 2001, also see Louda et al. 2003 for a review of 10 case studies with quantitative data), and the most obvious mechanisms that impact recipient communities appear to be extreme polyphagy in pair with the ability to infiltrate natural areas away from targeted agroecosystems. Interactions such as those existing between parasitoids and their hosts may also be impacted by the dynamics inherent to classical biological control. If they are able to, native non-specialist parasitoids may be displaced from their native hosts to an invasive one. However, the native community of parasitoids can be outcompeted from the exotic pest by the exotic parasitoid (Naranjo 2017) that has been chosen for its efficiency exploiting the exotic pest. Firstly, this could lead to local extinction of native populations of parasitoids, especially if the exotic parasitoid outcompetes them on the native host(s) as well (Bengtsson 1989). On the other hand, the introduction of an exotic parasitoid to control an exotic pest often leads to a displacement of the native community of parasitoids that have become associated with the exotic pest (Bennett 1993, Lynch and Thomas 2000, van Lenteren et al. 2006). This happens logically when the introduced parasitoid is specialized on the exotic pest and is a superior competitor or more adapted to find and exploit the pest than its native counterparts (Naranjo 2017). The resulting displacement might only be a step backwards, bringing the system back to the previous pattern of host-parasitoid dynamics (before the pest invaded the area), or a novel state might emerge, depending on the resilience of the native species. However, the temporal dynamics and spatial variability of these processes remain poorly understood and empirical data are greatly lacking at this point with, to our knowledge, no reports of such non-intentional effect in the context of biological control. Therefore, here we use successful classical biological control of an invasive pest as a framework to properly investigate how these two subsequent invasions impact the structure of native communities.

The Asian chestnut gall wasp *Dryocosmus kuriphilus* (Hymenoptera: Cynipidae), native to China, was accidentally introduced in Italy in 2002 (Brussino et al. 2002) and is now distributed throughout Italy and other European countries (EPPO, 2014). *D. kuriphilus* is a specialist attacking only chestnut trees. In absence of competitors (Bernardo et al. 2013) and specialized antagonists, D. *kuriphilus* was able to proliferate quickly and massively. Therefore, it became a trophic subsidy for several native parasitoids previously associated to gall wasps from other plants/trees (Matosevic and Melika 2013, Panzavolta et al. 2013, Francati et al. 2015, Noyes 2019). In response to damage observed on chestnut production and also apiculture, classical biological control programs were quickly implemented in newly infested countries. *Torymus sinensis* was chosen as a biological control agent due to its high specificity of parasitism (Quacchia et al. 2014, Ferracini et al. 2017) and its previous effective control of the target pest outside Europe (Gyoutoku and Uemura 1985, Cooper and Rieske 2007, 2011). In France, *T. sinensis* has been proven established with fast and significant impacts on the targeted pest in the subsequent years (Borowiec et al. 2018). This thus led to the quite unique opportunity to investigate how local communities evolve with regard to the deprivation of their trophic subsidy whereas most scientific work usually studies the recruitment of native parasitoids by the exotic biological control agents and its impact on food webs (Henneman and Memmott 2001, Eveleigh et al. 2007, Barber et al. 2008, Girardoz et al. 2008, David et al. 2017).

## Methodology

### Biological control introductions

In France, first isolated spots of *Dryocosmus kuriphilus* were observed from 2005 close to the Italian border but its pervasive presence in South of France was only patent from 2010. *Torymus sinensis* was introduced in France between 2011 and 2014, on a total of 59 sites (chestnut orchards) separated by at least 4 km. In each site, the monitoring of native parasitoids started one year prior to *T. sinensis* release. The introductions covered a wide geographical area (920 km from North to South, 1 030 km from East to West) in metropolitan France including Corsica. Two propagule sizes (100 and 1 000 individuals) of *T. sinensis* were introduced in separated sites but *T. sinensis* established itself in all sites whatever the initial propagule size was (see Borowiec et al. 2018 for more details). For this study we kept only the 26 sites for which at least five consecutive years of monitoring were available (Figure 1).

**Figure 1.**
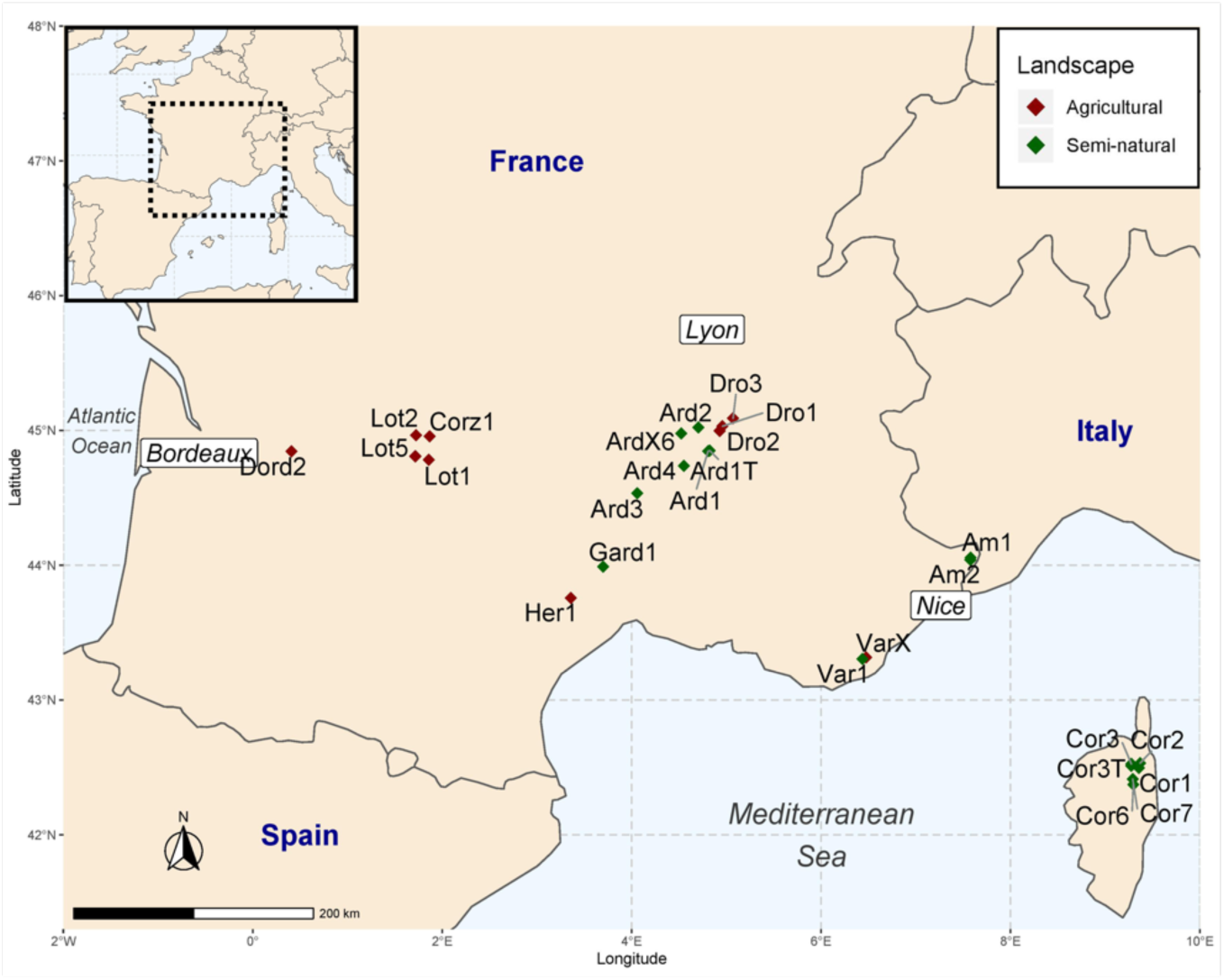
Map of the survey. Red points correspond to sites within an agricultural landscape (mainly apple orchards) whereas green points correspond to sites within a semi-natural landscape (mainly forests). Torymus sinensis was released in 2009 (Am1, Am2), in 2011 (Ard1, Cor1, Cor2, Cor3, Dro1, Dro2, Dro3, Var1) and in 2012 (Ard1T, Ard2, Ard3, Ard4, ArdX6, Cor3T, Cor6, Cor7, Corz1, Dord2, Gard1, Her1, Lot1, Lot2, Lot5, VarX).

### Sampling of insect communities associated with chestnut galls

#### Estimation of *D. kuriphilus* levels of infestation

Ten chestnut trees per site per year were sampled and, for each of them, ten totally random twigs were selected at human height and inspected for galls as explained in Table 1. The number of galls in a twig can therefore be zero. From these, the infestation levels of *D. kuriphilus* were estimated, by combining information on the mean percentage of buds with at least one gall and on the mean number of galls per bud as shown in Table 1. Because of the difficulty of this task (geographical cover, meteorological contingencies, staff’s availability and skills), infestations were finally not available for some sites and/or dates.

**Table 1.**
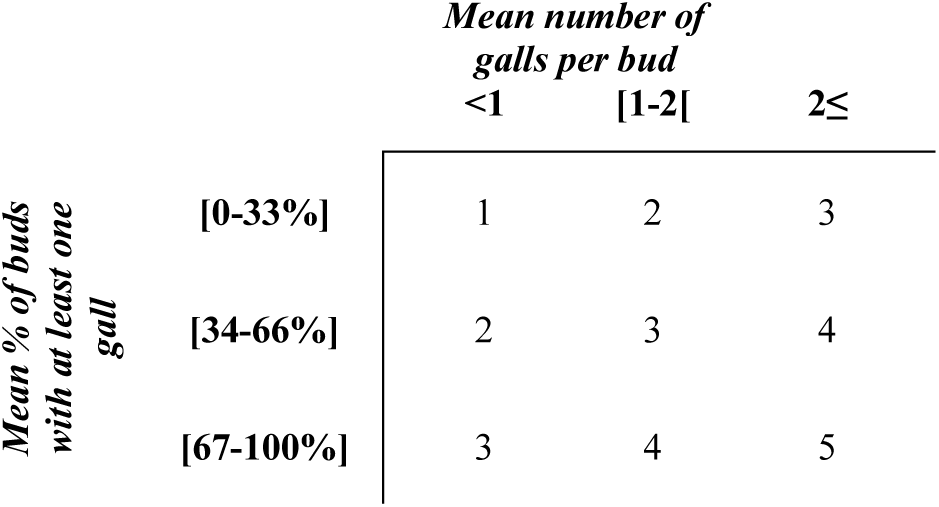
Table showing how the classes of infestation by D. kuriphilus were determined. Classes are created from 1 to 5 depending on the mean number of galls per bud and the mean percentage of buds with at least on gall. The higher these features are, the higher the infestation is estimated.

#### Diversity and abundance of associated parasitoids

As detailed in Borowiec et al. 2018, both the exotic *Torymus sinensis* and the native parasitoids (see Table 2) were counted from winter dry galls which are easily distinguished as they are brown and dry. Our work focuses on these galls and we deliberately omitted fresh spring galls for logistical reasons. In France, the Asian chestnut gall wasp is the only gall wasp on chestnut trees, therefore confusion with other species was impossible. We collected 2 000 to 5 000 galls per site during the two first years and then only 500 to 2 000. Galls were gathered on several trees. Once collected, galls were put in hermetic boxes placed outdoors from January to October so that parasitoids could develop and emerge in natural conditions at our laboratory (located at Sophia Antipolis, France). Each box referred to one site and contained 500 galls which constitutes a good compromise between the number of galls and the ability of parasitoids to emerge and get into the collecting vial placed at the extremity of the boxes. All emerged insects were collected and then stored in 96% ethanol and kept at −22°C. In addition to the exotic and ubiquitous *T. sinensis*, nine main native parasitoids were identified (Table 2). All identifications were based on morphological characters. *Eupelmus* species were identified using the latest descriptions of the *Eupelmus urozonus* complex (Al Khatib et al. 2014, 2016). Other species were identified by using an unpublished key to chalcidoid parasitoids in oak cynipid galls (Askew and Thuroczy, unpublished). DNA barcoding was used for a representative set of each morphologically-described genus and/or species to ascertain their identifications (see Molecular analyses section below).

**Table 2.** List of native parasitoids with their known host range. * This species may be a complex of cryptic parasitoid species but all specialized on Cynipidae. References: (1): Askew, R. R. and Thuroczy, (unpublished)– (2 Al Khatib, F. et al. (2014) (2016) – (3): Murakami et al. (1994) – (4): Noyes (2019).

| Native species | Host range | Refs |
| --- | --- | --- |
| *Aulogymnus* spp. | Hymenoptera :<br>Cynipidae | 1 |
| *Eupelmus azureus* | Hymenoptera :<br>Cynipidae | 2 |
| *Eupelmus kiefferi* | Coleoptera<br>Diptera<br>Hemiptera<br>Hymenoptera<br>Lepidoptera | 2 |
| *Eupelmus urozonus* | Diptera<br>Hymenoptera<br>Neuroptera | 2 |
| *Eurytoma setigera* | Hymenoptera :<br>Cynipidae : Cynipinae | 1, 3,<br>4 |
| *Megastigmus dorsalis* * | Hymenoptera :<br>Cynipidae | 1 |
| *Mesopolobus sericeus* | Hymenoptera :<br>Cynipidae : Cynipinae :<br>Cynipini | 1 |
| *Torymus auratus* | Hymenoptera :<br>Cynipidae : Cynipinae | 1, 4 |
| *Sycophila biguttata* | Hymenoptera :<br>Cynipidae | 1 |

### Molecular analyses

The DNA extraction was performed using commercial kits (Zygem PIN0500 or Quick extract Lucigen) according to the manufacturers’ recommendations in a total volume of 30µL, without crushing the insect. PCR targeted a small portion of the mitochondrial gene Cytochrome Oxidase I (COI), the standard barcode region (Hebert et al. 2003). We thus used the primers LCO 1490 (5’-GTCAACAAATCATAAAGATATTGG-3’) and HCO 2198 (5’-TAAACTTCAGGGTGACCAAAAAATCA-3’) (Folmer et al. 1994) or the related degenerated primers HCO_PUC (5’-TAAACTTCWGGRTGWCCAAARAAATCA-3’) and LCO_PUC (5’-TTTCAACWAATCATAAAGATATTGG-3’). When unsuccessful of PCR products or sequencing reaction, the primers COI pF2 (5’-ACCWGTAATRATAGGDGGDTTTGGDAA-3’) and COI 2437d (5’-GCTARTCATCTAAAWAYTTTAATWCCWG-3’) (developed by Simon et al. 1994 and modified by Kaartinen et al. 2010) were tried. The PCR conditions were as follows: 95°C for 5min, followed by 40 cycles of (i) 95°C for 30s, (ii) 48°C for 90sec, and 72°C for 1min with a final extension at 60°C for 30min. PCR products were shipped to Genewiz (Radolfzell, Germany) for their Sanger sequencing. The obtained sequences were checked and compared to either reliable sequences in GenBank (*Eupelmus* species – see Al Khatib et al. 2014, 2016), either to unpublished ones (see Appendix). Molecular analyses were carried out with MEGA-X software.

### Statistical analyses

#### Native parasitoid species co-occurrence analyses

To assess the patterns of co-occurrence between parasitoid species, and their evolution over time, we used the C-score (Stone and Roberts 1990) from each annual matrix of presence-absence of the nine native species. As *T. sinensis* was always present and here we only consider species occurrences (presence/absence), it was excluded of the analysis.

The C-score is a measure of average pairwise species segregation. It measures the mean number of checkerboard units between all pairs of species in a data matrix. The number of checkerboard units for each pair of species *i* and *k* is calculated as follows:

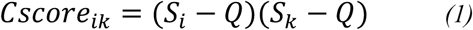

where Q is the number of shared sites, S_i_ and S_k_ are the number of sites in which species *i* and *k* are respectively found. In equation (1) the C-score will be equal to zero if species *i* and *k* share all sites. Conversely, C-score will be equal to one if species species *i* and *k* are never found together. The overall C-score for the community is then calculated as follows:

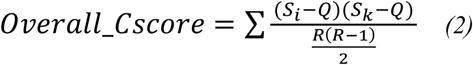

where R is the number of rows (=species) in the matrix (Stone and Roberts 1990, Gotelli 2000). When compared to other co-occurrence indices such as CHECKER (Diamond 1975), V-ratio (Robson 1972, Schluter 1984) and COMBO (Pielou and Pielou 1968), C-score has the smallest probability of type I and II errors (Gotelli 2000). However, because the value of the C-score depends on the frequency of occurrence of the species, inter-annual comparisons cannot be performed directly. We thus used the co-occurrence null model from the EcoSimR package (Gotelli 2015) of R (R Development Core Team 2018) to create null assemblages based on our observed presence-absence species matrices. This was done by randomizing (by transposing sub-matrices) species occurrences but keeping row and columns totals fixed (Gotelli 2000). Thus, differences between sites are maintained, making this method appropriate to detect patterns of species interactions (Gotelli 2000). Each randomization produces one matrix in which a ‘simulated’ C-score is calculated. Such randomization is replicated ten thousand times. The significance of the observed C-scores was computed as the proportion of simulated values equal to or more extreme than the observed C-score value.

In order to graphically compare each year, all c-score values were normalized by using:

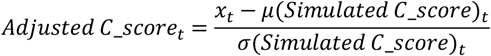

where *x* takes the values of observed and simulated C-score and t refers to the year after release of *T. sinensis* (from 1 to 5). A low value of C-score is indicative of an aggregative pattern, while a high value is indicative of an exclusion pattern.

#### Native parasitoids community structure

We described the community structure each year after the release of *T. sinensis* by using the R package ‘pheatmap’ (Raivo 2019). We created clustered heatmaps with the ‘pheatmap’ function to visualize how communities of native parasitoids are structured during the survey. Sites were clustered depending on their native parasitoid absolute abundance using aggregative clustering. As a distance measure between clusters x and y we used the Euclidean distance which is calculated as follows:

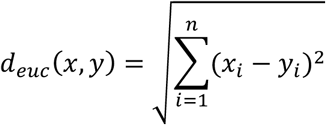

As a linkage function, we chose the complete linkage which takes the maximum distance between the two clusters:

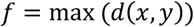

Aggregative clustering starts by computing a distance between every pair of units to be clustered and creating a distance matrix. The two clusters with the smallest value (e.g. A and B) are then merged into a new cluster (e.g. AB). The matrix is then recalculated and as we use the complete linkage the distance between the new cluster and the other clusters (e.g. C and D). The distance between AB and C will be equal to the maximum value between C and A and between C and B.

#### Landscape context

In an attempt to evaluate the potential role of habitat on the community structure during the last year of survey, we used a Principal Component Analysis (PCA) considering the abundances of each native species using the ‘FactoMineR’ package of R (Husson et al. 2019). More precisely, sites were plotted in the first dimension of the PCA depending on the abundance of native parasitoids in each of them. We then plotted the two main categories of habitat: (i) orchards located within an agricultural landscape with a poor amount of semi-natural habitat; (ii) orchards located within semi-natural habitats (mainly forested areas). The habitat categorization was made based on qualitative estimates first during field work and confirmed later on satellites views. Satellite views were necessary in order to confirm habitat type within the 1-km radius of field observations for all areas belonging to private owners, preventing our access and direct categorization of habitats.

To do so, we used the Open-source platform QGIS (https://qgis.org/en/site/) with the satellite view of the OpenLayers plugin. We also used the land registry layer of the French government land registry website (https://www.geoportail.gouv.fr/donnees/registre-parcellaire-graphique-rpg-2017).

We also tested with a generalized linear model whether the abundance of *M. sericeus* was significantly different in agricultural and semi-natural habitats. However, given the numerous zeros within the agricultural category, we settled with a generalized linear model testing whether the occurrence (modelled as binomial) of *M. sericeus* was significantly different between the two habitats.

## Results

### Control of *Dryocosmus kuriphilus* by *Torymus sinensis*

A fast increase of the *Torymus sinensis*’ relative frequency was observed during the 5 years of survey, 90% of galls being finally parasitized by *T. sinensis* (Figure 2). This was easily monitored as each gall can only contain one *T. sinensis* inside. In parallel, the infestation levels of *D. kuriphilus* decreased markedly, based on the sites in which infestation data was available for each year of the survey (Am1, Am2, Ard1, Gard2, Var1).

**Figure 2.**
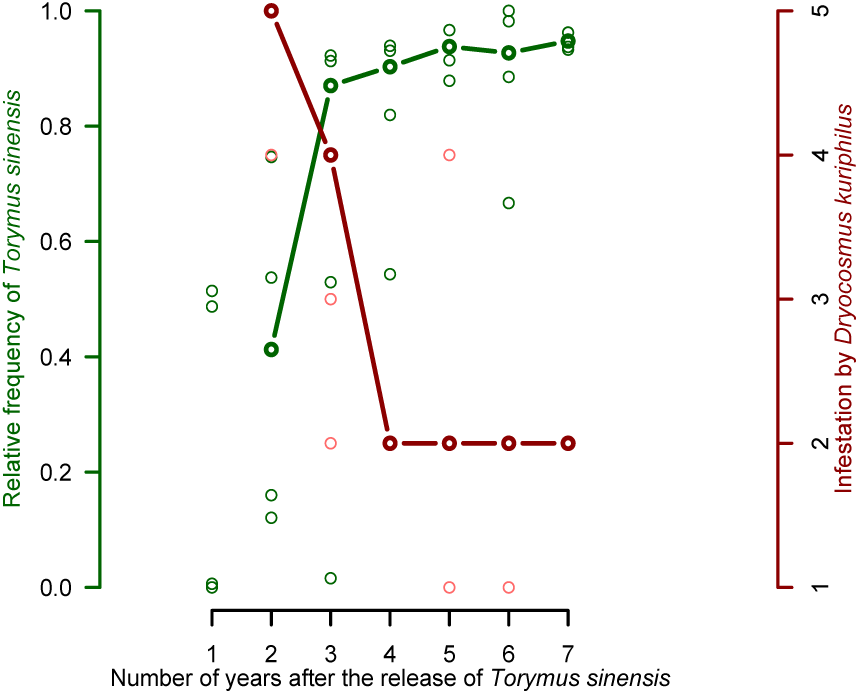
Infestation levels of D. kuriphilus and Relative frequency of T. sinensis in galls each year of the survey. Here are only shown the sites where the infestation was consistently measured for all years of the survey.

### Molecular-assisted species identification

The identification of individuals was realized in routine using morphological characters. The molecular characterization was however necessary for taxa in which a species’ complex is known (for instance, in the *Eupelmus* genus – see Al Khatib et al. 2014 and 2016) or for which few information is available. As shown in the Figure S2, the COI sequenced (between 550 and 612pb) were informative enough to distinguish closely related species, as in the *Sycophyla* and *Torymus* genera. For some taxa (*Aulogymnus arsames, Eurytoma setigera, Megastigmus dorsalis, Torymus affinis*), the within molecular diversity may suggest the presence of sister species and/or a marked intraspecific variability.

### Abundances and occurrences of native species

Overall, 71 494 specimens of *T. sinensis* and 12 016 specimens of native parasitoids were obtained from 284 425 galls from the 26 sites during the 5 years of the survey. Galls from all locations were monitored for emerging parasitoids at our laboratory.

In terms of abundance, the native species were ordered as follows: *Mesopolobus sericeus* (n= 3 792, 31.6%), *Eupelmus urozonus* (n= 2 069, 17.2%), *Megastigmus dorsalis* (n= 1 877, 15.6%), *Eupelmus azureus* (n= 1 752, 14.6%), *Eurytoma setigera* (n=586, 4.9%), *Eupelmus kiefferi* (n=491, 4.1%), *Aulogymnus* spp. (n=403, 3.4%), *Sycophyla biguttata* (n=116, 1%), *Torymus auratus* (n=19, 0.1%). Nine hundred and eleven (7.5%) individuals remained undetermined using both morphological and molecular identification and were thus discarded from the analysis.

In terms of occurrence, *Torymus sinensis* was observed in the 130 possible site-by-year combinations. In comparison, the results for native species were as follows: *Eupelmus urozonus* (n=111), *Eurytoma setigera* (n=86), *Eupelmus kiefferi* (n=74), *Megastigmus dorsalis* (n=59), *Mesopolobus sericeus* (n=50), *Eupelmus azureus* (n=49), *Aulogymnus* spp. (n=33) *Sycophyla biguttata* (n=26), *Torymus auratus* (n=7).

The mean abundances of all nine native parasitoids are given for each year in Figure S1 (*Supplemental Figure S1*). Overall, they peaked during the second and/or third years of the survey.

A potential pitfall with such survey could be to miss some other relevant species because of some insufficient sampling effort. However, we are quite confident that this is not the case insofar as, excepted for the rarest species (*T. auratus*), all other species were found at least common or even very common in some site-by-date combination, well above a potential detection threshold (Figure 3).

**Figure 3.**
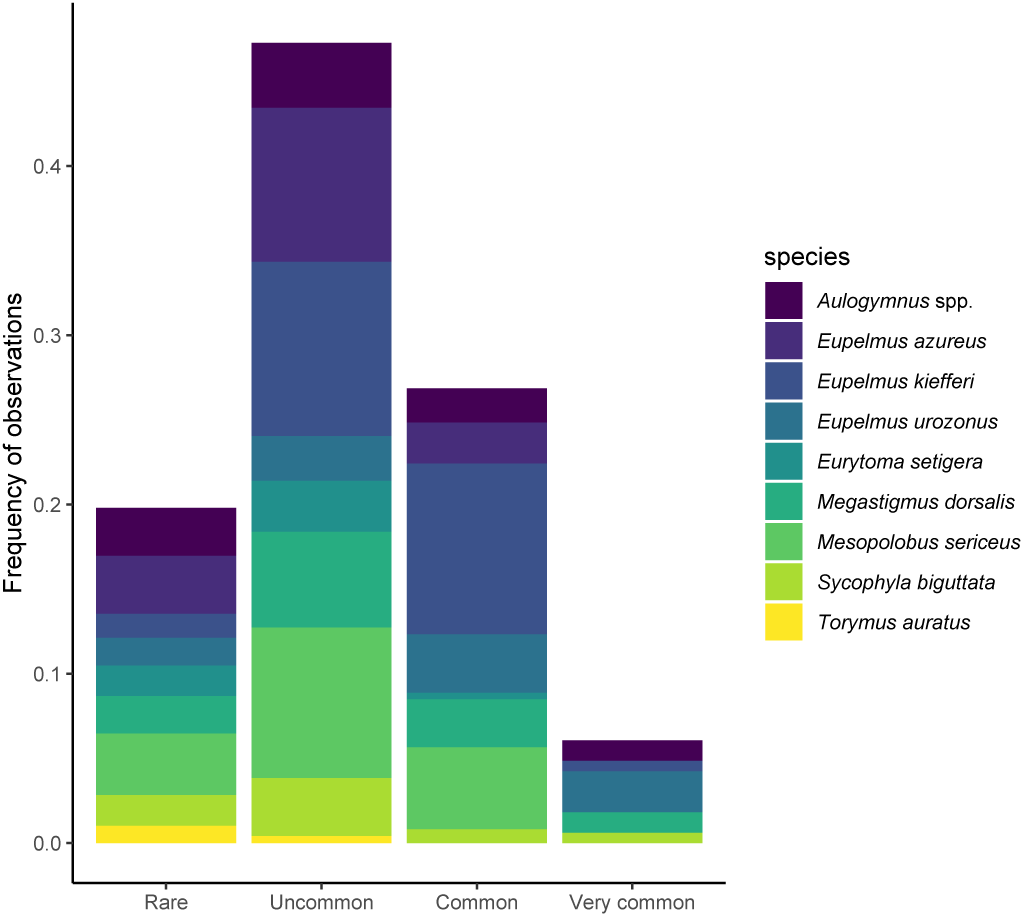
Frequencies of observations the different species of native parasitoids. Site-by-date-by-species combinations were sorted according to four classes of abundance: Rare (only 1 individual of the species), Uncommon (from 2 to 10 individuals), Common (from 11 to 100 individuals) and Very common (From 101 to max abundance sampled).

### Co-occurrence null model analyses

Starting the third year, native parasitoids species co-occurred less frequently than expected by chance with increasing odds from the first to the fifth year (Figure 4). Therefore, as years passed, there was a decreasing chance of observing co-occurrence of native parasitoids by sampling *D. kuriphilus* galls. More details about which pairs of species were co-occuring more frequently across the years can be extracted from the heatmaps made for each year of the survey (supplementary material Figure S2-S5).

**Figure 4.**
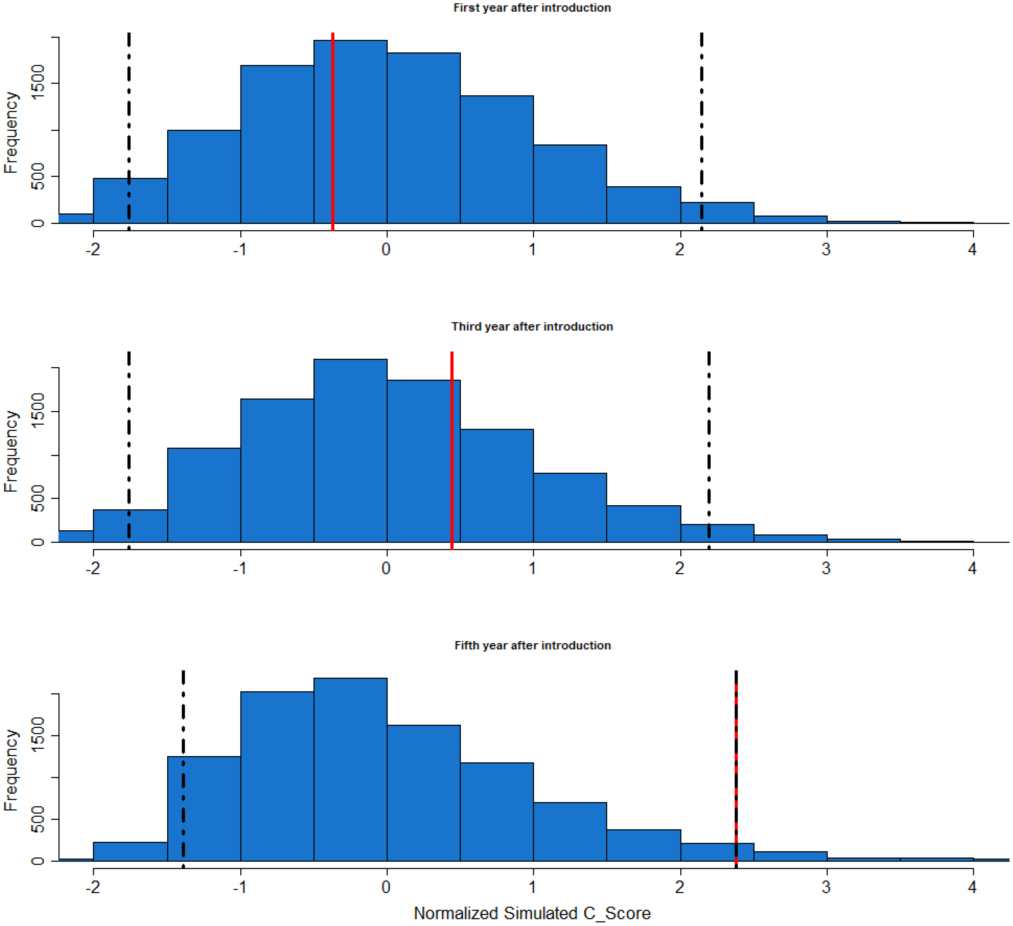
C-score values for the native community of parasitoids for the first (A), third (B) and fifth year (C) after the release of T. sinensis. Blue histogram represents the simulated values, the red bar represents the observed C-score and the dotted lines represent the 95% confidence interval.

### Native parasitoid community structure

During the fifth year of the experiment, we observed two different patterns among sites (Figure 5). In the first category (19 sites, top cluster), the community of native parasitoids was represented by just a few species occurring at low abundances. Furthermore, in a few sites (Lot5, Corz1, Dord2), not even a single native parasitoid was sampled. However, in the second category (7 sites, bottom cluster), the community was dominated by *Mesopolobus sericeus*, a specialized parasitoid of the Cynipini tribe (Table 2) that was not sampled anywhere the first year of the survey (supplementary material, Figure S2).

**Figure 5.**
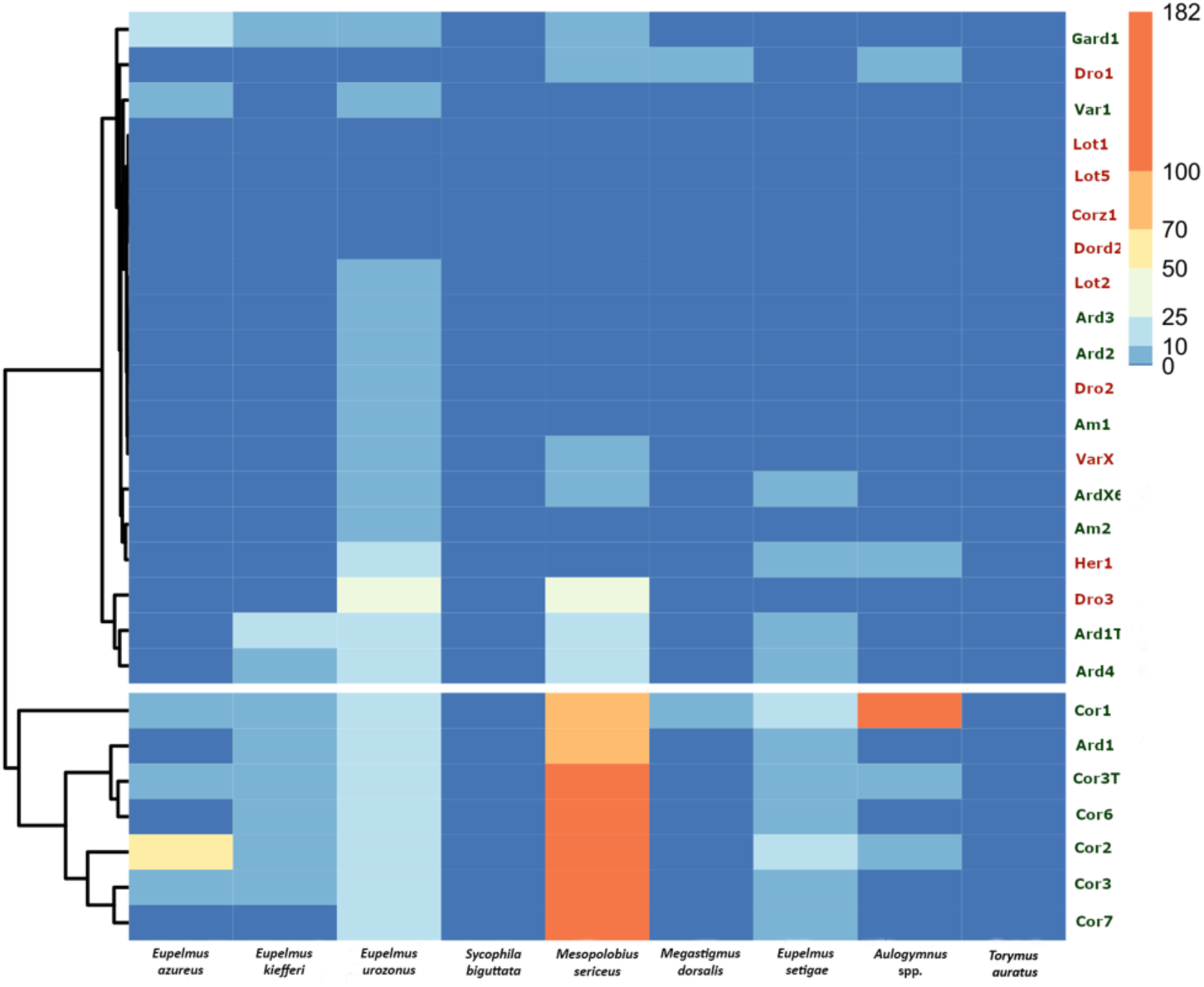
Heatmap representing the abundances of all native parasitoid species during the fifth year after the release of T. sinensis. Abundances are represented in log scale by a color gradient from blue to red. The colors of locations’ names refer to the type of habitat (red: agricultural habitat – green: semi-natural habitat). To help the reader, species names are ordered as follows (from left to right): Eupelmus azureus, Eupelmus kiefferi, Eupelmus urozonus, Sycophila biguttata, Mesopolobus sericeus, Megastigmus dorsalis, Eupelmus setigae, Aulogymnus spp. Torymus auratus.

### Landscape context

The Principal Component Analysis on native parasitoid abundances confirmed that, five years after the release of *T. sinensis*, communities were mostly structured by the local presence of *M. sericeus* (Figure 6A). The analysis of the projection of the different sites highlighted that the abundance of *M. sericeus* was correlated with the type of habitat, semi-natural orchards being more likely to host this particular species (Figure 6B). As the PCA suggested an effect driven mostly by the response of the species *M. sericeus*, we showed that this species was indeed occurring more frequently within semi-natural orchards (p.value=0.0396).

**Figure 6.**
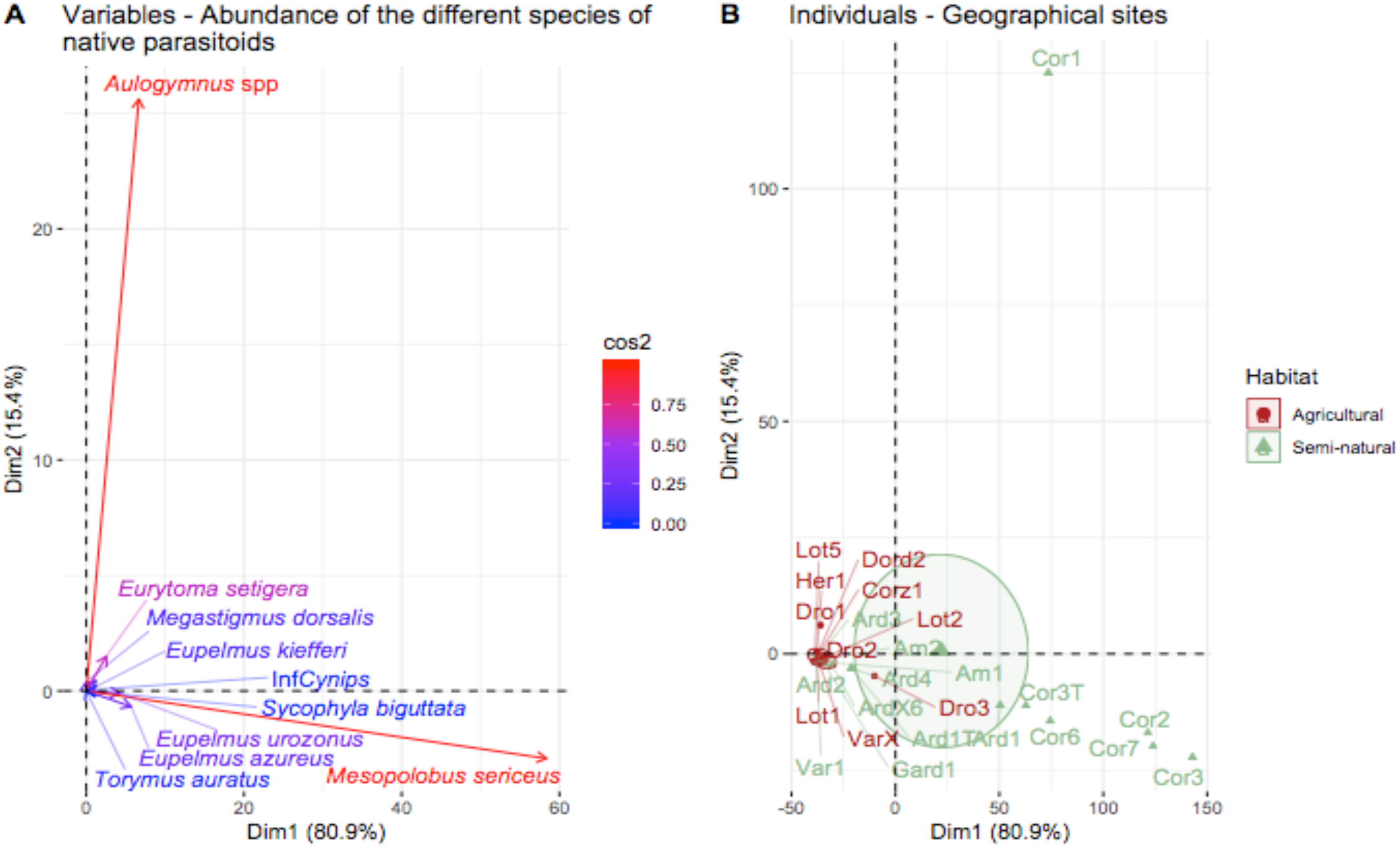
**A**: Variables correlation plot of the PCA built from the abundance of native parasitoids Colors represent the contribution of the different species to the structure of the variation within each axis. **B**: Projection of the different geographical sites on two principal component axis of the PCA. The colors discriminate the two main habitat: Agricultural (red) and Semi-natural (green).

To verify whether the strong pattern driven by *M. sericeus* might have obscured responses in other species, we also ran the analysis by excluding *M. sericeus*. We found that the native parasitoid community still evolves towards segregation, although it is only significant the fifth year of the survey (supplementary material Figure S6).

The clustering was significantly different with virtually no difference between sites. Here all sites (excepted Cor1) are grouped in the same big cluster (supplementary material Figure S7). Likewise, the PCA showed a weaker differentiation between our two types of habitat (supplementary material Figure S8).

## Discussion

Our work highlights the restructuration of native parasitoid communities following the successful control of *Dryocosmus kuriphilus* by the biological control agent *Torymus sinensis*. Our results suggest that communities evolved differently mostly depending on the landscape context of the sampling site.

The nine native parasitoids that appeared to use *D. kuriphilus* as a trophic subsidy in our survey are related to oak gall wasps, although their degree of generalism is highly variable (Table 2). Some species such as *Eupelmus kiefferi* and *E. urozonus* are indeed extremely polyphagous whereas, in contrast, *Mesopolobus sericeus* is substantially more specific, being specialized on only one tribe of Cynipids (Noyes 2019). The main result of the co-occurrence model analyses is the increasing exclusive competition through time (Figure 4). This means that the native community underwent a significant restructuration, which was correlated in time with the rarefaction of *D. kuriphilus*. The first parasitoids actually exploiting *D. kuriphilus* were *E. urozonus, E. azureus* and *M. dorsalis* (Figure S1), those species combining (i) a wide host range, (ii) a known affinity with several Cynipidae and (iii) pre-existing resources in such kinds of habitats (Al khatib et al. 2014, Al khatib et al. 2016). They were thus therefore likely to shift relatively easily on the new invasive host (Cornell and Hawkins 1993, Hawkins 2005). However, in the following years, *Mesopolobus sericeus*, which was not detected during the first year of survey (Figure S1), was the sole species able to markedly increase in abundance and to persist on this trophic subsidy (Figure 5). And this, despite the fact that it was not detected once at the beginning of the survey in any site (Figure S1). It seems to be the same for *Aulogymnus* spp. except that these are much less abundant and include all species from the *Aulogymnus* genus. The examples of native species displaced by invasive species are numerous (e.g. Rowles and O’Dowd 2007, Bohn et al 2008, Inoue et al 2008, Sebastian et al 2015) therefore we are not surprised to observe such outcome for most of our native species, which are outcompeted by the introduced specialist *T. sinensis*. Only *Mesopolobus sericeus* appears to be able to coexist at significant levels with *Torymus sinensis* on *Dryocosmus kuriphilus*. Furthermore, the most generalist parasitoids of our system probably exhibit a switching behavior (Murdoch 1969). This behavior refers to a situation where a predator (or here a parasitoid) exhibits adaptive, flexible choices between all available preys (or hosts). This choice entails positively frequency-dependent predation. In our study abundance of other hosts are unknown. It is therefore possible that *D. kuriphilus* eventually becomes rarer than other hosts, forcing generalists to switch to more abundant hosts (Pelletier 2000). However, switching preys can have stabilizing or destabilizing effects, even possibly causing extinction of the predator (Van Leeuwen et al 2007).

Increasing landscape complexity (mostly defined as the increase of the proportion of semi-natural habitat) is generally associated with increases in natural enemy abundance and/or diversity (Bianchi et al. 2006, Schmidt et al. 2008, Gardiner et al. 2009a, 2009b). In our study, sites enclosed within large amounts of semi-natural habitats contained the most diverse and abundant native parasitoid communities (Figure 5). In particular, the persistence of *M. sericeus*, the most specialist native parasitoid, was modulated by the local environment (Figure 6B). Indeed, *M. sericeus* was the native parasitoid that seemed to be the best at exploiting *D. kuriphilus*, although only within semi-natural habitat. Among the seven sites showing a marked domination of *M. sericeus* (Figure 5), six of which are located in the island of Corsica (Figure 5). Islands are home to plant and animal communities with relatively little diversification, simplified trophic webs and high rates of endemism (Williamson 1981, Chapuis et al. 1995). In addition to their smaller size, these characteristics (Cassey 2003) partially explain that oscillations (or perturbations) within the resident community are more incline to destructive outcomes such as extinctions (Elton 1958). Nonetheless, five continental sites (Figure 5: ArdX6, Dro3, Ard1T, Ard4, Ard1) also exhibit a slightly less marked but similar increase of *M. sericeus* (Figure 5), four of them being in semi-natural landscapes (Figure 1). We thus think that the final dominance of *M. sericeus* towards other native species is rather explained by differences in the landscape rather than a “mainland *versus* island” dichotomy. In fact, most of the known hosts of *M. sericeus* are oak gall wasps (Noyes 2019), oak trees being rarer in agricultural landscapes than in semi-natural ones. Large populations of *M. sericeus* acting like sources for the colonization of chestnut orchards are consequently more likely to be sustained in this latter habitat.

In classical biological control, the ecological impacts of a biological control agent are usually explained by the use of non-target hosts/preys (Louda et al. 2003). Although *Torymus sinensis* has shown a slight host range expansion it appeared to be with minimal impact and no effect expected on distribution and abundance of non-target hosts (Ferracini et al. 2017). Thus, its impact on native competitors is mediated by its successful control of the shared host.

Our sampling effort was strong enough that we can be confident we did not miss any relevant species. Even if a single individual of a given species was sampled relatively frequently (about 20% of all data), all species except the rarest one were also sampled at high density in some cases. This implies that no species in our analyses was so rare that the probability to have completely missed another species with comparable abundance by chance is close to zero. With regard to the rarest species, *Torymus auratus*, it is already known to be rare in winter dry galls but more abundant in spring fresh galls, so that our sampling method might have underestimated its abundance (Kos et al. 2015, Ferracini et al. 2018). Furthermore, species rankings according to abundance and occurrence are not the same, which confirms that our sampling method is robust enough to accurately detect species even when their abundance is low.

We need to point out that although our study contains insightful information on how our native parasitoid community structure evolves, species dynamics we observed the last year of the survey are not fixed but quite the opposite. Species dynamics are most probably still evolving towards a, yet unknown, state of equilibrium. We are still not in measure to predict with certainty what will happen when *D. kuriphilus* will become even rarer. Maybe *T. sinensis* will remain the dominant species or maybe because of the presence of its native hosts, *M. sericeus* will outperform *T. sinensis* at least in semi-natural habitats. Furthermore, although we evidenced a successful host range expansion (by newly including *T. sinensis* into their diet) from the majority of these native parasitoids, nothing is known about how the populations dynamics evolved on their native hosts.

In conclusion, classical biological control offers an exciting frame to investigate real-time population dynamics during invasive processes. Yet, the opportunities remain rare because of various reasons including (i) the quite high rate of establishment’s failure, (ii) the temporal frame required for the observation of significant patterns and (iii) the lack of funding for post-release surveys. With regard to this context, the deliberate introduction of *T. sinensis* against *D. kuriphilus* in France thus was a quite unique opportunity. Our work sheds a new light on how the “boom-and-bust” (here defining the situation in which a period of great prosperity is abruptly followed by one of decline) dynamics of an invasive pest can impact the structure of native communities of potential antagonists. Our results evidence a site-specific scenario where a sole native species, *M. sericeus*, dominates the native community on the trophic subsidy and is able to co-exist with the exotic and specialized competitor, *T. sinensis. M. sericeus* is now able to exploit both the native gall wasps and *D. kuriphilus*. This extended host range may have lasting impacts on *T. sinensis* populations, all the more so *D. kuriphilus* will reach a low density at a global scale. In turn, the rarefaction of *D. kuriphilus* and the competition with *M. sericeus* might constrain *T. sinensis* to exploit new hosts. Therefore, it would be of particular interest to study the long-term evolution of these two species as the ideal expected outcome of classical biological control is the everlasting control of the host with no unintentional negative impact on the recipient community. Another open perspective of this work is to analyze how the structure of native parasitoids evolves within the oak gall wasp’s community.

## Data accessibility

Data are available online: https://doi.org/10.5281/zenodo.3929233

## Code accessibility

Script and codes are available online: https://doi.org/10.5281/zenodo.3952462

## Acknowledgements

We thank the University of Torino for fruitful collaborations and the Interreg Alcotra project ‘Sauvegarde de l’ecosystème châtaigne’ for initial funding. We also thank the ‘Syndicat National des Producteurs de châtaignes’ as well as several extension services (CTIFL, INVENIO, FREDON, Chambres d’Agriculture) for their help in releasing T. sinensis and participating to post-release surveys. Finally, we would like to thank Jean-Claude Malausa who initiated this project, and the ISA-team ‘Research and Development in Biological Control’ for their contribution to intensive field sampling. The biological control program against the chestnut gall wasp in France was granted by the National Program ECOPHYTO (‘CYNIPS’, 2011–2014; ‘CYNIPS2’, 2016–2017) and by the Plant Health and Environment Division of INRAE (2015).

Version 6 of this preprint has been peer-reviewed and recommended by Peer Community In Zoology (https://doi.org/10.24072/pci.zool.100004).

## Conflicts of interest disclosure

The authors of this preprint declare that they have no financial conflict of interest with the content of this article. Elodie Vercken is a recommender of PCI EvolBiol and PCI Ecology.

## Notes

### Competing Interest Statement

The authors have declared no competing interest.

