## Supplementary_material for "The open bar is closed: restructuration of a native parasitoid community following successful control of an invasive pest"

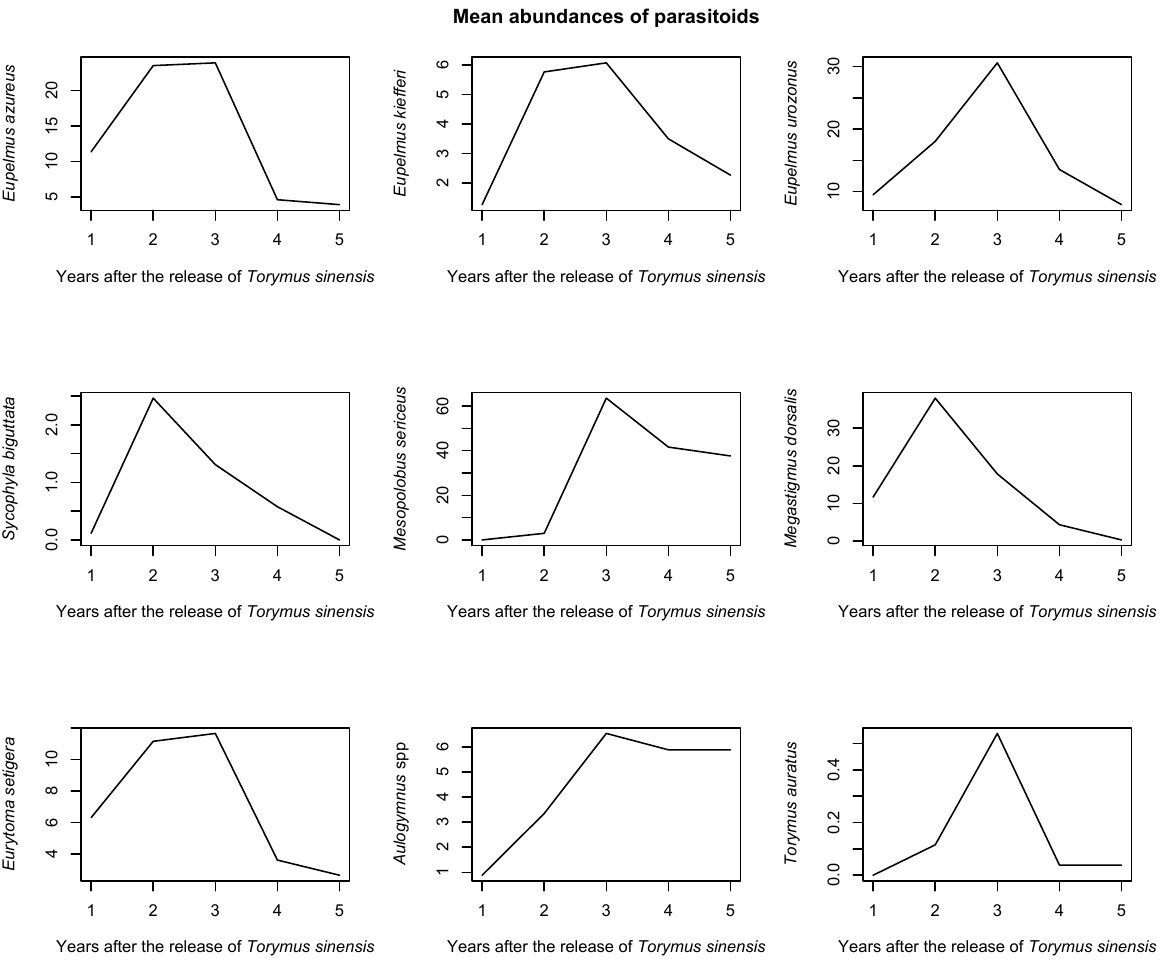

Figure S1 - Mean abundances of all native species during the five years of the study

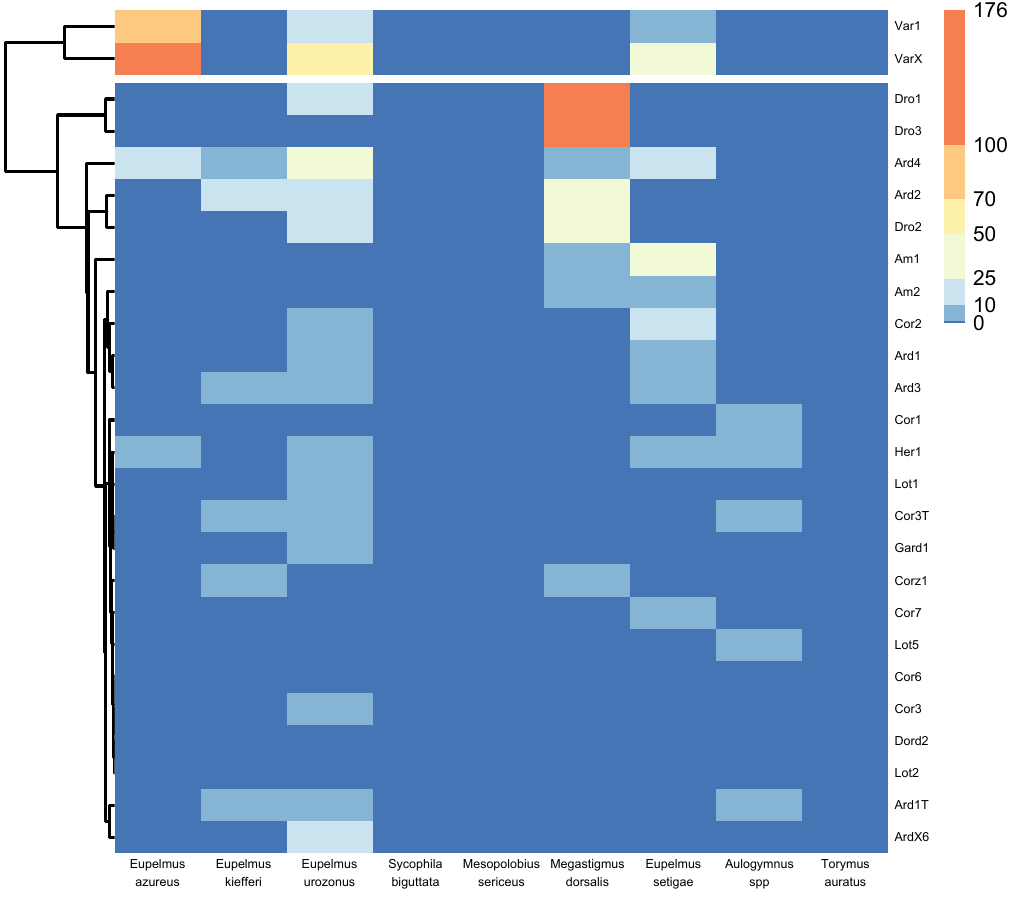

Figure S2 - Heatmap for the first year of the survey

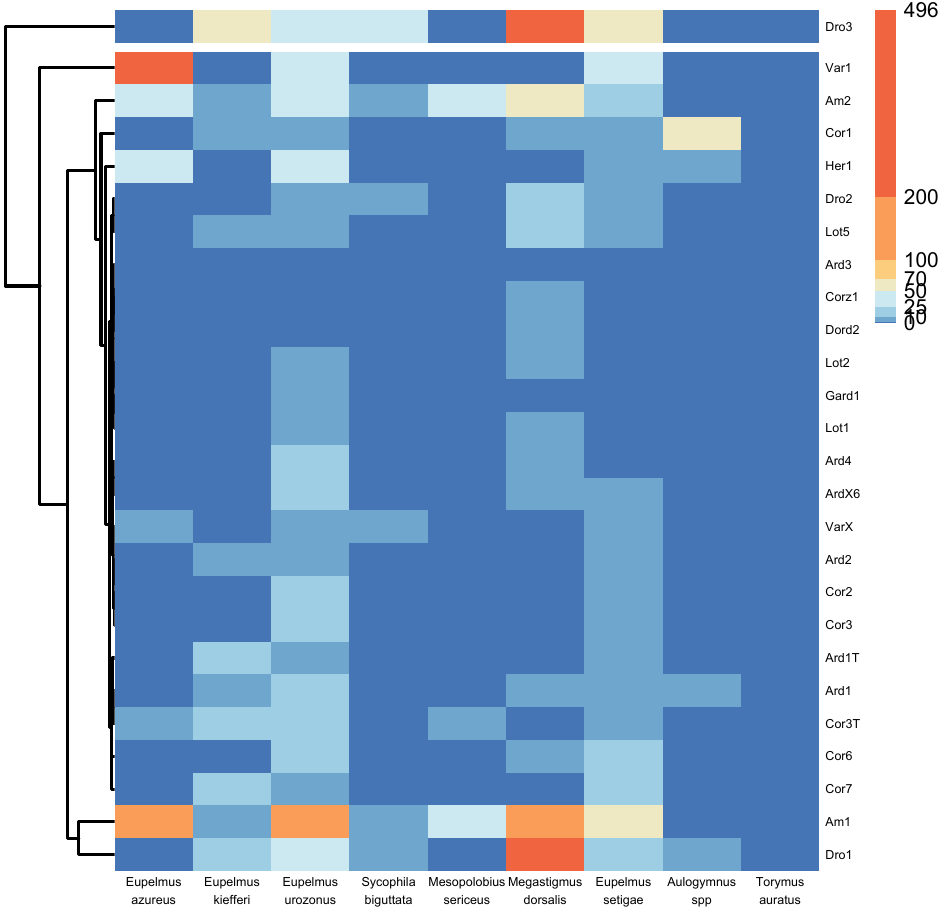

Figure S3 - Heatmap for the second year of the survey

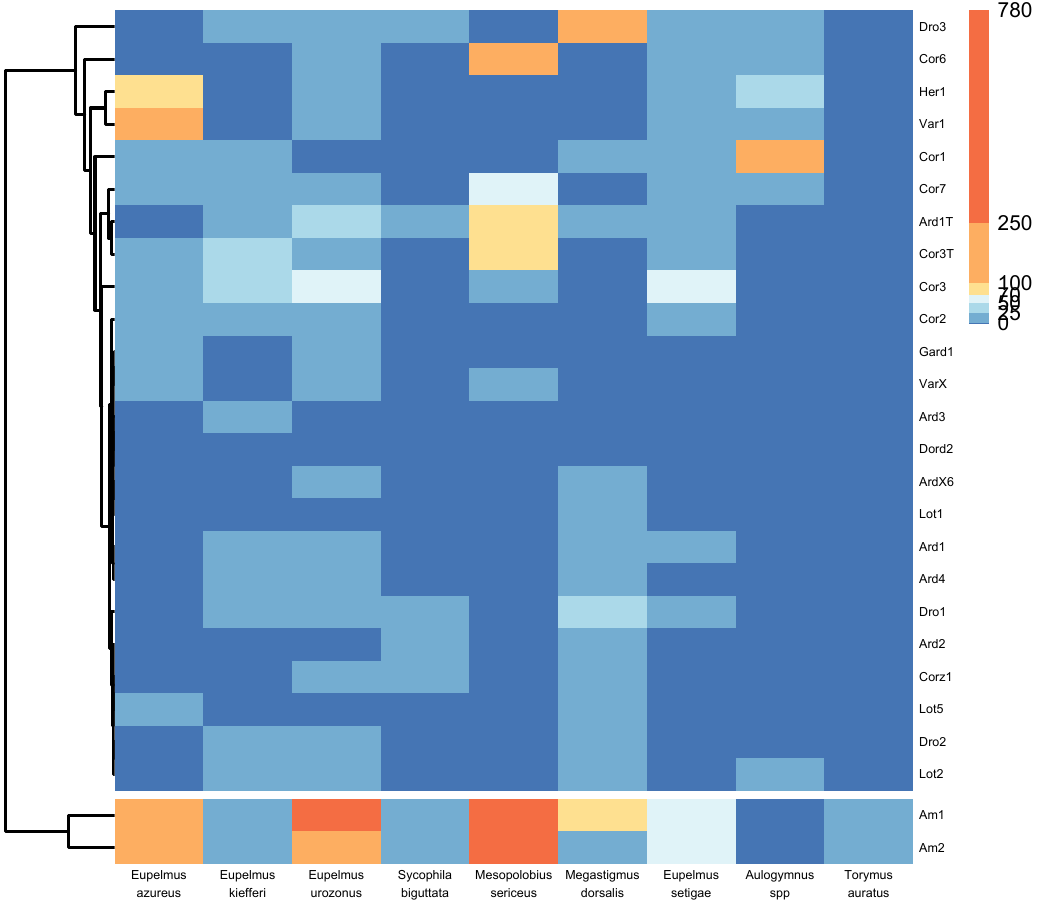

Figure S4 - Heatmap for the third year of the survey

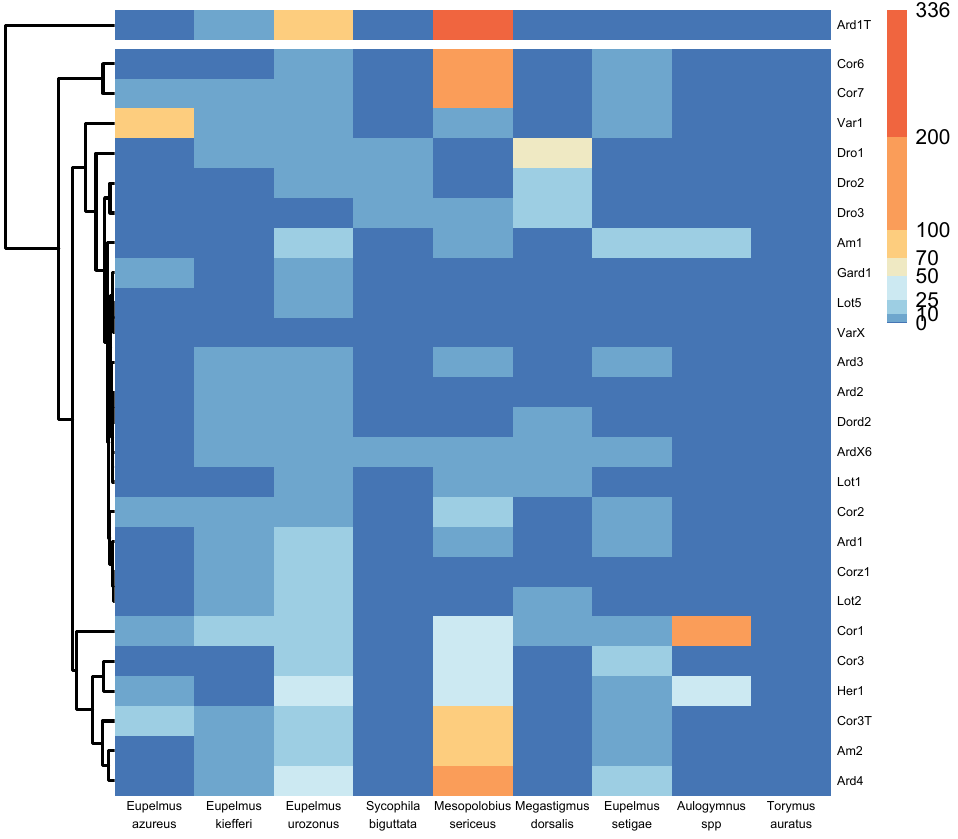

Figure S5 - Heatmap for the fourth year of the survey

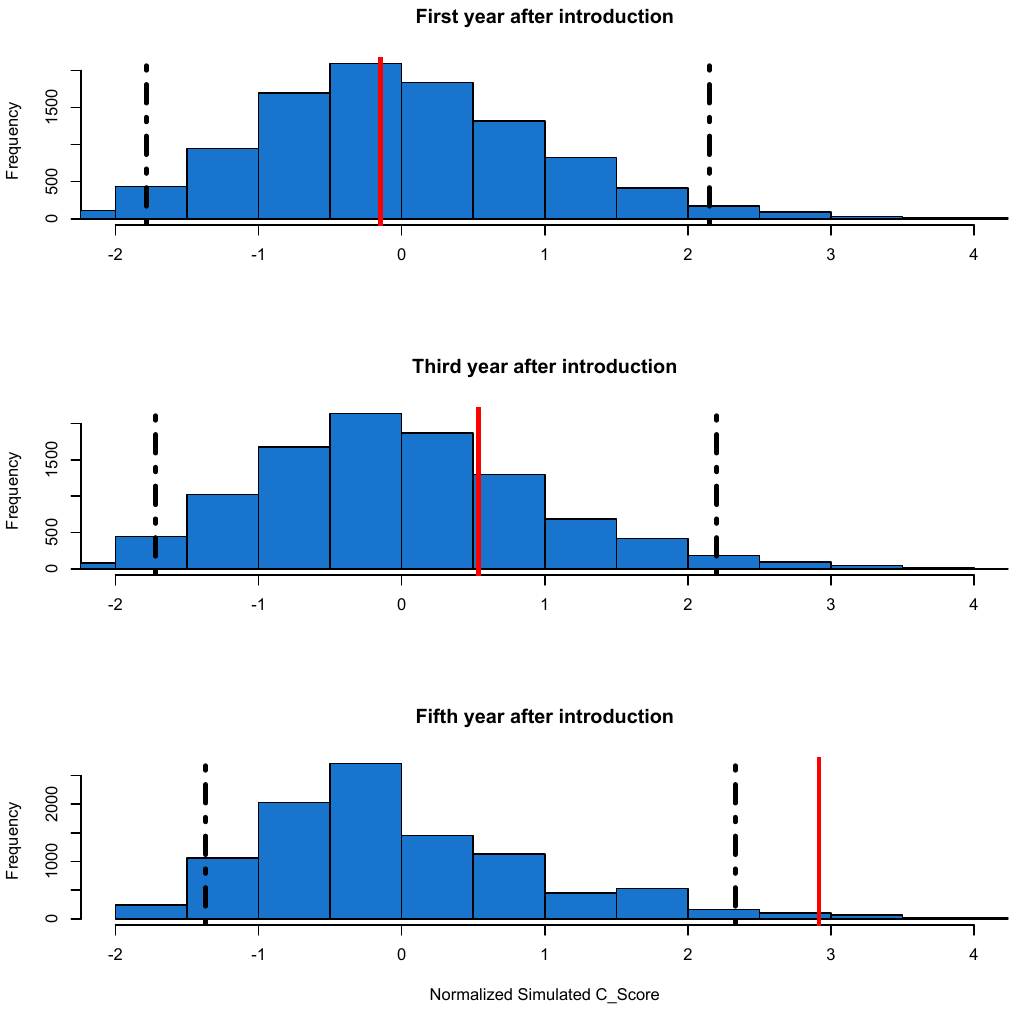

Figure S6 - Co-occurrence analysis excluding Mesopolobus sericeus

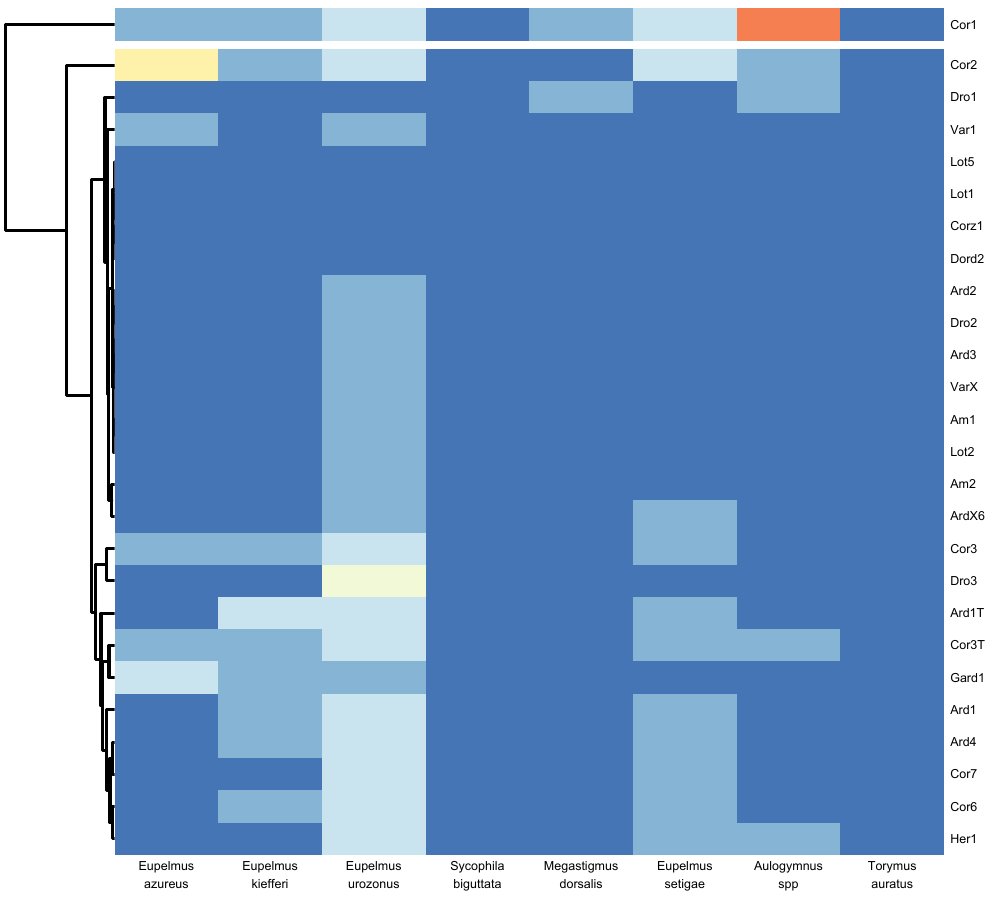

Figure S7 - Heatmap for the fifth year of the survey, exlcuding Mesopolobus sericeus

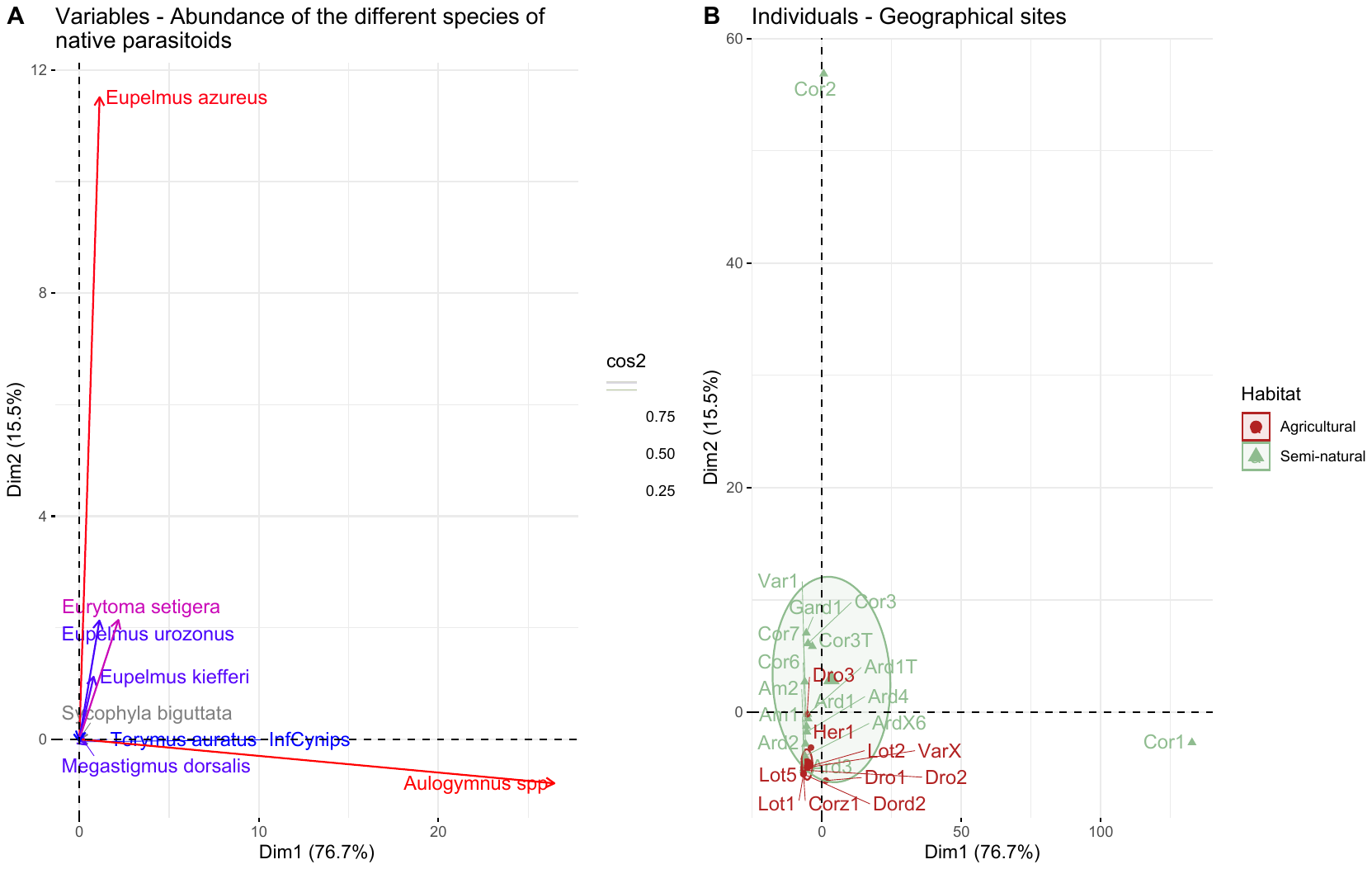

Figure S8 - PCA for the fifth year of the survey excluding Mesopolobus sericeus

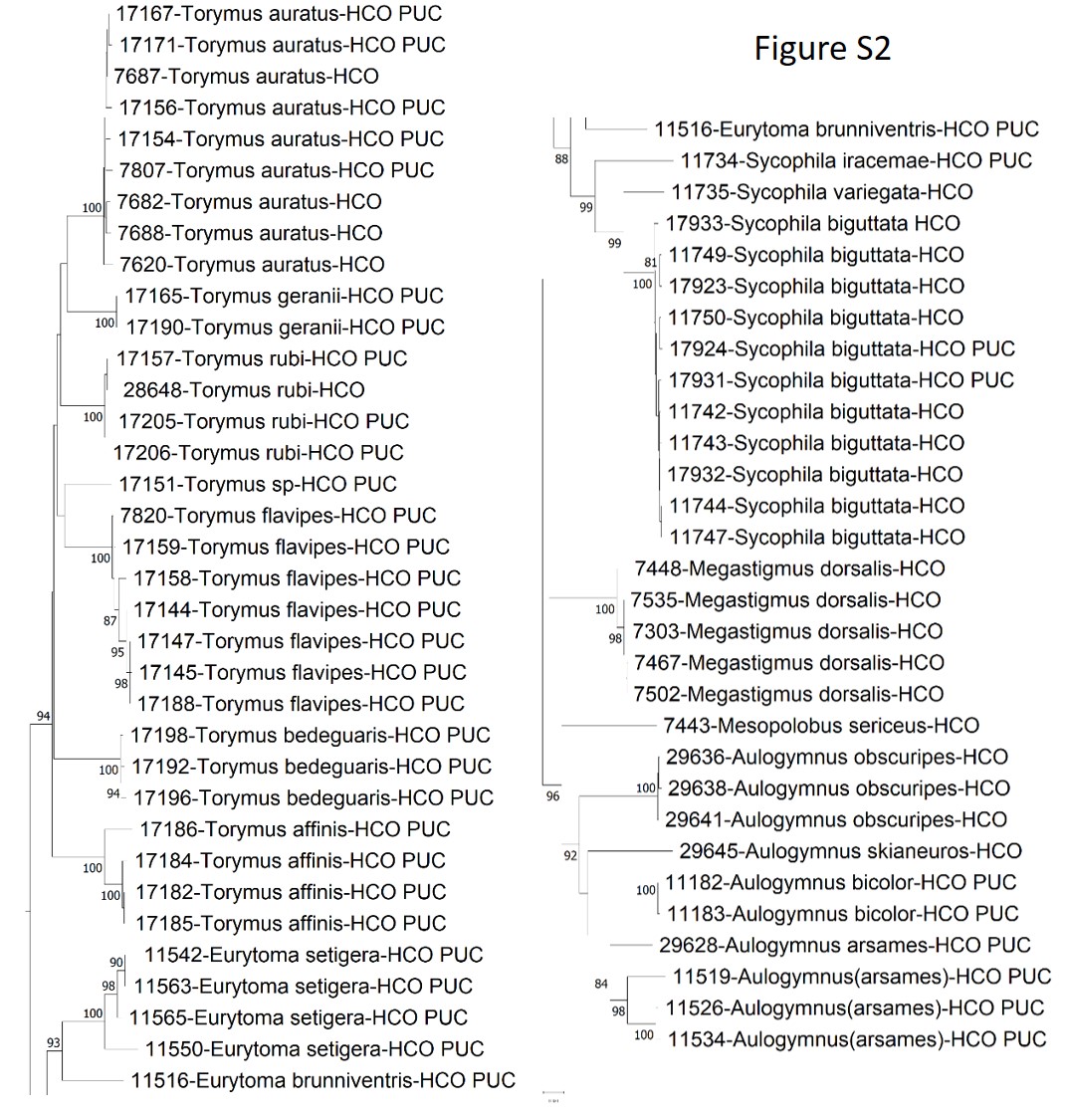

Figure S 9 - Neighbour joining tree obtained from the COI sequences detailed in Appendix 1. This was obtained using MEGA-X with the following parameters: Kimura 2 parameters distance, pairwise deletion (sequences between 550 and 612pb) and 500 replicates for bootstrapping. The labels are organized as follows (from left to right): (1) numeric code: internal code allowing to obtain additional information, (2) genus and species names based on morphological criteria (uncertain species name in brackets), (3) Primer used. The individuals were extracted from a more comprehensive dataset encompassing several hundreds of parasitoids from Dryocosmus kuriphilus and other gallwasps. Those selected here illustrate the haplotypic diversity within each taxa.

**Appendix : Unreleased sequences of *Aulogymnus*, *Eurytoma*, *Megastigmus*, *Mesopolobus*, *Sycophila* and *Torymus*.**

These sequences were extracted from a more comprehensive dataset involving several hundreds of parasitoids reared from *Dryocosmus* *kuriphilus* or from other gallwasps. The 64 specimens selected here illustrated the observed haplotypic diversity. Each of these specimens were also carefully morphologically characterized in particular by Nicolas Borowiec (co-author) and by Ionela-Madalina Viciriuc (co-author) For some *Aulogymnus* specimens, the identification was uncertain and the species name was indicated in bracket.

>11182-Aulogymnus_bicolor-HCO_PUC

TATTTTTGGAATATGAGCTGGAATTTTGGGATTATCAATAAGATTAATAATTTGATTAGAGTTAGGAAATCCTGGATCCTTGATTGGAAATGATCAAATTTATAATTCAATTGTTTCAACCCATGCATTTACAATAATTTTTTTTTTTGTTATACCAGTAATAATAGGAGGATTTGGAAATTATTTGATTCCTTTAATATTAGGGACTCCAGATATAGCTTTTCCTCGAATAAATAATATAAGATTTTGATTACTTCCACCAAGATTAATATTATTAATTTCAAGAATATTTATTGGTTCAGGGACTGGGACAGGATGAACTGTTTATCCACCATTATCTTCAAATTTAGGGCATAGAGGTCCTTCGGTTGATTTATCAATTTTTTCTTTACATATTGCTGGTATTTCTTCCATTATAGGTTCAATTAATTTTATTAGAACAATTTTAAATATAAAAAATTATAAAATAGAAAATATTTCATTATTTTCATGATCAATATTATTAACTGCAATTTTATTACTCTTATCTCTTCCTGTTTTAGCTGGTGCTA

>11183-Aulogymnus_bicolor-HCO_PUC

TATTTTTGGAATATGAGCTGGAATTTTGGGATTATCAATAAGATTAATAATTTGATTAGAGTTAGGAAATCCTGGATCCTTGATTGGAAATGATCAAATTTATAATTCAATTGTTTCAACCCCTGCATTTACAATAATTTTTTTTTTTGTTATACCAGTAATAATAGGAGGATTTGGAAATTATTTGATTCCTTTAATATTAGGGACTCCAGATATAGCTTTTCCTCGAATAAATAATATAAGATTTTGATTACTTCCACCAAGATTAATATTATTAATTTCAAGAATATTTATTGGTTCAGGGACTGGGACAGGATGAACTGTTTATCCACCATTATCTTCAAATTTAGGGCATAGAGGTCCTTCGGTTGATTTATCAATTTTTTCTTTACATATTGCTGGTATTTCTTCCATTATAGGTTCAATTAATTTTATTAGAACAATTTTAAATATAAAAAATTATAAAATAGAAAATATTTCATTATTTTCATGATCAATATTATTAACTGCAATTTTATTACTCTTATCTCTTCCTGTTTTAGCTGGTGCTA

>11516-Eurytoma_brunniventris-HCO_PUC

AAAGATATTGGAATTTTATATTTTATTTTTGGAATATGAGCAGGAATTTTAGGTCTTTCATTGAGAATAATTATTTGAATAGAATTAGGAAGACCTGGATCATTAATTGGTAATGATCAAATTTATAATTCAATTGTTACTACTCATGCTTTTGTTATAATTTTTTTTTTTGTAATACCTGTAATGATAGGGGGATTTGGAAATTTTTTAATTCCATTAATTTTAGGAATTCCTGATATAGCATTTCCTCGAATAAATAATATAAGTTTTTGATTATTAGTACCTAGATTAATATTATTAATTTCTAGAATATTTATTGGAAATGGTACTGGTACAGGTTGAACAGTATATCCTCCTCTTTCAGGAAATTTATCTCATGGAGGTCCTTCAGTAGATTTATCTATTTTTTCTTTGCATATTGCAGGGGTTAGATCAATTATAGGATCTATTAATTTTATTTCAACTATTTTAAATATAAAAATTTATAAAATAGAATTAATTCCTTTATTTGCTTGAGCAATATTATTAACTACAATTTTATTACTTTTATCTTTACCAGTGTTAGCTGGAGCTATTACTATATTATTATTTGATCGAAATTTAAA

>11519-Aulogymnus(arsames)-HCO_PUC

TATTTTTGGAATATGAGCTGGAATTTTAGGGTTGTCAATAAGATTAATAATTTGTTTAGAATTAGGAAATCCTGGATCCTTAATTGGTAATGACCAGATTTATAATTTTATTGTCCCTACTCATGCATTTACTATAATTTTTTTTTTTGTTATACCAGTAATAATAGGGGGATTTGGAAATTATTTAATTCCTTTAATGTTGGGTGTCCCAGATATAGCTTTCCCTCGAATAAATAATATAAGCTTTTGATTACTTCCTCCAAGATTAATATTATTAATTTCTAGAATATTTATTGGATCTGGAACGGGTACGGGATGAACAGTTTACCCTCCTTTGTCTTCTAATTTAGGACATAGAGGCCCATCAGTAGATTTATCAATTTTTTCTCTTCATATTGCTGGTGCATCTTCAATTATAGGGTCTATTAATTTTATCAGAACAATTTTAAATATAAAAAATTATAAGATAGAAAATATTTCTTTATTTTCATGATCGATATTATTAACTGCAATTTTATTACTTTTATCTCTTCCTGTATTAGCTGGTGCAA

>11526-Aulogymnus(arsames)-HCO_PUC

TATTTTTGGAATGTGAGCTGGGATTTTAGGGTTATCAATAAGATTAATAATTTGTTTAGAATTAGGAAATCCTGGATCTTTAATTGGTAATGATCAGATTTATAATTTTATTGTTACTACTCATGCATTTACTATAATTTTTTTTTTTGTGATACCAGTAATAATAGGAGGATTTGGAAATTATTTAATTCCTTTGATATTAGGCGTTCCAGATATAGCTTTCCCTCGAATAAATAATATAAGATTTTGACTTCTTCCTCCAAGATTAATATTATTAATTTCTAGAATATTTATTGGGTCTGGGACGGGTACAGGATGAACAGTTTATCCTCCTTTGTCATCTAATTTAGGACATAGGGGACCATCAGTAGATTTATCAATTTTTTCTCTTCATATTGCTGGGATATCATCAATTATAGGATCTATTAATTTTATTAGTACAATTTTAAATATAAAAAATTATAAGATAGAAAATATTTCTTTATTTTCATGATCAATATTATTAACTGCAATTTTATTACTTTTGTCTCTTCCTGTACTAGCTGGAGCAA

>11534-Aulogymnus(arsames)-HCO_PUC

TATTTTTGGAATGTGAGCTGGAATTTTAGGGTTATCAATAAGATTAATAATTTGTTTAGAATTAGGAAATCCCGGATCTTTAATTGGTAATGATCAGATTTATAATTTTATTGTTACTACTCATGCATTTACTATAATTTTTTTTTTTGTGATACCAGTAATAATAGGAGGATTTGGAAATTATTTAATTCCTTTGATATTAGGGGTTCCAGATATAGCTTTCCCTCGAATAAATAATATAAGATTTTGACTTCTTCCTCCAAGATTAATATTATTAATTTCTAGAATATTTATTGGGTCTGGGACGGGTACAGGATGAACAGTTTATCCTCCTTTGTCATCTAATTTAGGACATAGGGGACCATCAGTAGATTTATCAATTTTTTCTCTTCATATTGCTGGGATATCATCAATTATAGGATCTATTAATTTTATTAGTACAATTTTAAATATAAAAAATTATAAGATAGAAAATATTTCTTTATTTTCATGATCAATATTATTAACTGCAATTTTACTACTTTTGTCTCTTCCTGTACTAGCTGGAGCAA

>11542-Eurytoma_setigera-HCO_PUC

AAAGATATTGTAATTTTATATTTTATTTTTGGAATATGAGCGGGAATTTTAGGGTTATCTTTAAGAATAATTATTTGAATGGAATTAGGTAATCCTGGTTCTTTAATTGGAAATGATCAAGTTTATAATTTTATTGTTACAACTCATGCTTTTATTATAATTTTTTTTTTTGTAATACCTGTAATAATAGGGGGGTTTGGAAATTTTTTAGTTCCAATAATTTTAGGAATTCCTGATATAGCATTTCCTCGGATAAATAATATGAGTTTTTGATTATTAATTCCTAGAATTACTTTATTAATTTTTAGAATATTTATTGGTAGGGGAACTGGGACAGGTTGAACAGTTTATCCTCCTTTATCAGGGAATTTATTTCATGGGGGGCCTTCAGTAGATTTATCGATTTTTTCTTTACATGTTGCTGGAGTTAGATCAATTATAGGATCAATTAATTTTATTAATACAGTTTTAAATATAAAAATTTATAAAAGTGAATTAATTCCTTTGTTTGCTTGATCAATATTATTGTCTACTATTTTATTATTATTATTTTTTCCAGTTTTAACAGGTGCAATTACCATAATATTATATGATCGAAATTTAAA

>11550-Eurytoma_setigera-HCO_PUC

AAAGATATTGGAATTTTATATTTTATCTTTGGGATATGAGCGGGAGTTTTAGGATTATCTTTAAGAATAATTATTTGAATAGAATTAGGTAATCCTGGTTCTTTAATTGGAAACGATCAAGTTTATAATTTTATCGTTACAACTCATGCTTTTATTATAATTTTTTTTTTTGTGATACCTGTAATAATAGGGGGTTTTGGAAATTTTTTGGTTCCAATAATTTTAGGGATTCCTGATATAGCATTTCCTCGAATAAATAATATAAGTTTTTGATTATTAATTCCTAGAATTACTTTACTAATTTCTAGAATATTTATTGGAAGGGGGACTGGGACAGGTTGAACAGTTTATCCTCCTTTATCAGGAAATTTATCTCATGGGGGTCCTTCAGTAGATTTATCTATTTTTTCTTTACATGTTGCGGGGGTCAGATCAATTATAGGATCAATTAATTTTATTAATACAGTTTTAAATATAAAAATTTATAAAATTGAATTAATTCCTTTATTTGCTTGATCAATGTTATTGACAACTATTTTACTATTATTATTTTGTCCAGTTTTAGCAGGTGCAATTACCATATTATTATATGATCGAAATTTAAA

>11563-Eurytoma_setigera-HCO_PUC

AAAGATATTGGAATTTTATATTTTATTTTTGGAATATGAGCGGGAATTTTAGGGTTATCTTTAAGAATAATTATTTGAATGGAATTAGGTAATCCTGGTTCTTTAATTGGAAATGATCAAGTTTATAATTTTATTGTTACAACTCATGCTTTTATTATAATTTTTTTTTTTGTAATACCTGTAATAATAGGGGGGTTTGGAAATTTTTTAGTTCCAATAATTTTAGGAGTTCCTGATATAGCATTTCCTCGGATAAATAATATGAGTTTTTGATTATTAATTCCTAGAATTACTTTATTAATTTTTAGAATATTTATTGGTAGGGGAACTGGGACAGGTTGAACAGTTTATCCTCCTTTATCAGGGAATTTATTTCATGGGGGGCCTTCAGTAGATTTATCGATTTTTTCTTTACATGTTGCTGGAGTTAGATCAATTATAGGATCAATTAATTTTATTACTACAGTTTTAAATATAAAAATTTATAAAATTGAATTAATTCCTTTGTTTGCTTGATCAATATTATTAACTACTATTTTATTATTATTATTTTTTCCAGTTTTAGCAGGTGCAATTACCATAAAATTATATGATCGAAATTTAAA

>11565-Eurytoma_setigera-HCO_PUC

AAAGATATTGGAATTTTATATTTTATTTTTGGAATATGAGCGGGAATTTTAGGGTTATCTTTAAGAATAATTATTTGAATGGAATTAGGTAATCCTGGTTCTTTAATTGGAAATGATCAAGTTTATAATTTTATCGTTACAACTCATGCTTTTATTATAATTTTTTTTTTTGTAATACCTGTAATAATAGGGGGTTTTGGAAATTTTTTAGTCCCAATAATTTTAGGAATTCCTGATATAGCATTTCCTCGGATAAATAATATGAGTTTTTGATTATTAATTCCTAGAATTACTTTATTAATTTCTAGAATATTTATTGGTAGGGGAACTGGGACAGGTTGAACAGTTTATCCTCCTTTATCAGGGAATTTATCTCATGGGGGGCCTTCAGTAGATTTATCGATTTTTTCTTTACATGTTGCTGGAGTTAGATCAATTATAGGATCAATTAATTTTATTAATACAGTTTTAAATATAAAAATTTATAAAATTGAATTAATTCCTTTGTTTGCTTGATCAATATTATTAACTACTATTTTATTATTATTATCTTTTCCAGTTTTAGCAGGTGCAATTACCATATTATTATATGATCGAAATTTAAA

>11734-Sycophila_iracemae-HCO_PUC

AAAGATATTGGTATTTTATATTTAATTTTTGGGATATGGTTTGGAATTTTAGGTTTATCATTAAGAATATTGATTCGATTGGAATTAGGTAATCCTGGTTCTTTAATTGGAAATGATCAGATTTATAATTCAATTGTTACAGCTCATGCTTTTATCATAATTTTTTTTTTTGTAATGCCTGTTATAATAGGTGGATTTGGAAATTATTTGATTCCTTTAATTTTAGGAATTCCTGATATGGCCTTTCCTCGAATAAATAATATAAGTTTTTGATTATTAATTCCAAGATTAATTTTGTTAATTTCTAGAATATTTGTAGGTTCTGGAACAGGAACTGGTTGGACAGTTTACCCTCCTTTATCAGGTAATTTGTCTCATGGTGGTCCTTCAGTAGATTTATCTATTTTTTCTCTCCATTTAGCGGGAGTAAGATCTATTATAGGTTCTGTAAATTTTATTTCTACAATTATTAATATAAAAATTAGAAAGATTGAGTTGATTCCTTTATTTGCTTGATCTATATTATTAACTACTATTTTGTTACTATTGTCTCTTCCTGTTTTAGCAGGGGCTATTACTATATTATTATTTGATCGGAATTTGAA

>11735-Sycophila_variegata-HCO

AAAGATATTGGAGTTTTATATTTAATTTTTGGGATATGAAGAGGGGTTTTAGGTTTATCTTTAAGTATACTAATTCGTTTAGAATTAGGAAATCCTGGCTCTTTAATTGGTAACGATCAGATTTATAATTTTATTGTTACTGCTCATGCTTTTATTATAATTTTTTTTTTTGTTATACCTGTGATGATAGGAGGGTTTGGAAATTATTTAATTCCTTTAATCTTAGGAATTCCAGATATAGCTTTTCCTCGAATGAATAATATAAGATTTTGGTTATTAATTCCTAGATTATTTTTGTTACTTTCTAGAATATTTGTTGGTTCAGGTACTGGAACTGGTTGGACTGTTTATCCTCCTTTATCTGGGAATTTATCGCATGGAGGTCCATCAGTTGATTTATCAATTTTTTCACTTCATTTAGCTGGAGTTAGTTCAATTATAGGTTCAGTAAATTTTATTTCTACTATTTTGAATATAAAGATTTTTAAAATTGAGTTAATTCCTTTATTTGCTTGATCTATATTATTAACTACAATTTTATTACTTTTGTCATTACCAGTTTTAGCTGGAGCTATTACTATACTACTTTTTGATCGAAATTTGAA

>11742-Sycophila_biguttata-HCO

AAAGATATTGGAGTTTTATATTTAATTTTTGGGATATGAAGTGGAGTTTTAGGTTTATCTTTAAGAATATTAATTCGGTTAGAATTAGGGAATCCTGGTTCTTTAATTGGTAATGATCAAATTTATAATTCAATTGTCACTGCTCATGCTTTTATTATAATTTTTTTTTTTGTTATACCTGTTATGATAGGGGGATTTGGAAATTATTTAATTCCTTTAATTTTAGGAATTCCTGATATAGCTTTTCCTCGAATAAATAATATAAGATTTTGATTATTAATTCCTAGATTATTTTTATTACTTTCAAGTATATTTGTAGGTTCGGGAACTGGAACTGGATGGACCGTTTATCCACCTTTATCTGGAAATTTATCTCATGGTGGACCATCAGTTGATTTATCTATTTTTTCTCTTCATTTAGCTGGTATTAGTTCAATTATAGGTTCAGTTAATTTTATTTCTACAATTTTGAATATAAAAATTTTTAAAATTGAGTTAATTCCTTTATTTGCTTGATCAATATTATTAACTACGATTTTGTTACTTTTGTCATTACCAGTTTTAGCTGGAGCTATTACTATATTACTTTTTGACCGAAATTTAAA

>11743-Sycophila_biguttata-HCO

AAAGATATTGGAGTTTTATATTTAATTTTTGGGATATGAAGTGGAGTTTTAGGTTTATCTTTAAGAATATTAATTCGGTTAGAATTAGGAAACCCTGGTTCTTTAATTGGTAATGATCAAATTTATAATTCAATTGTCACTGCTCATGCTTTTATTATAATTTTTTTTTTTGTTATACCTGTTATGATAGGGGGATTTGGAAATTATTTAATTCCTTTAATTTTAGGAATTCCTGATATAGCTTTTCCTCGAATAAATAATATAAGATTTTGATTATTAATTCCTAGATTATTTTTATTACTTTCAAGTATATTTGTAGGTTCGGGAACTGGAACTGGATGGACCGTTTATCCACCTTTATCTGGAAATTTATCTCATGGTGGACCATCAGTTGATTTATCTATTTTTTCTCTTCATTTAGCTGGTATTAGTTCAATTATAGGTTCAGTTAATTTTATTTCTACAATTTTGAATATAAAAATTTTTAAAATTGAGTTAATTCCTTTATTTGCTTGATCAATATTATTAACTACGATTTTGTTACTTTTGTCATTACCAGTTTTAGCTGGAGCTATTACTATATTACTTTTTGACCGAAATTTAAA

>11744-Sycophila_biguttata-HCO

AAAGATATTGGAGTTTTATATTTAATTTTTGGGATATGAAGTGGAGTTTTAGGTTTATCTTTAAGAATATTAATTCGGTTAGAATTAGGAAATCCTGGTTCTTTAATTGGTAATGATCAAATTTATAATTCAATTGTCACTGCTCATGCTTTTATTATAATTTTTTTTTTTGTTATACCTGTTATGATAGGGGGATTTGGAAATTATTTAATTCCTTTAATTTTAGGAATTCCTGATATAGCTTTTCCTCGAATAAATAATATAAGATTTTGATTATTAATTCCTAGATTATTTTTATTACTTTCAAGTATATTTGTAGGTTCGGGAACTGGAACTGGATGGACCGTTTATCCACCTTTATCTGGAAATTTATCTCATGGTGGACCATCAGTTGATTTATCTATTTTTTCTCTTCATTTAGCTGGTATTAGTTCAATTATAGGTTCAGTTAATTTTATTTCTACAATTTTGAATATAAAAATTTTTAAAATTGAGTTAATCCCTTTATTTGCTTGATCAATATTATTAACTACGATTTTGTTACTTTTGTCATTACCAGTTTTAGCTGGAGCTATTACTATATTACTTTTTGACCGAAATTTAAA

>11747-Sycophila_biguttata-HCO

AAAGATATTGGAGTTTTATATTTAATTTTTGGGATATGAAGTGGAGTTTTAGGTTTATCTTTAAGAATATTAATTCGGTTAGAATTAGGAAATCCTGGTTCTTTAATTGGTAATGATCAAATTTATAATTCAATTGTCACTGCTCATGCTTTTATTATAATTTTTTTTTTTGTTATACCTGTTATGATAGGGGGATTTGGAAATTATTTAATTCCTTTAATTTTAGGAATTCCTGATATAGCTTTTCCTCGAATAAATAATATAAGATTTTGATTATTAATTCCTAGATTATTTTTATTACTTTCAAGTATATTTGTAGGTTCGGGAACTGGAACTGGATGGACCGTTTATCCGCCTTTATCTGGAAATTTATCTCATGGTGGACCATCAGTTGATTTATCTATTTTTTCTCTTCATTTAGCTGGTATTAGTTCAATTATAGGTTCAGTTAATTTTATTTCTACAATTTTGAATATAAAAATTTTTAAAATTGAGTTAATCCCTTTATTTGCTTGATCAATATTATTAACTACGATTTTGTTACTTTTGTCATTACCAGTTTTAGCTGGAGCTATTACTATATTACTTTTTGACCGAAATTTAAA

>11749-Sycophila_biguttata-HCO

AAAGATATTGGAGTTTTATATTTAATTTTTGGGATATGAAGTGGAGTTTTAGGTTTATCTTTAAGAATATTAATTCGGTTAGAATTAGGAAATCCTGGTTCTTTAATTGGTAATGATCAAATTTATAATTCAATTGTCACTGCTCATGCTTTTATTATAATTTTTTTTTTTGTTATACCTGTTATGATAGGGGGATTTGGAAATTATTTAATTCCTTTAATTTTAGGAATTCCTGATATAGCTTTTCCTCGAATAAATAATATAAGATTTTGATTATTAATTCCTAGATTATTTTTATTACTTTCAAGTATATTTGTAGGTTCGGGAACTGGGACTGGGTGGACCGTTTATCCACCTTTATCTGGGAATTTATCTCATGGTGGACCATCAGTTGATTTATCTATTTTTTCTCTTCATTTAGCTGGTATTAGTTCAATTATAGGTTCAGTCAATTTTATTTCTACAATTTTGAATATAAAAATTTTTAAAATTGAGTTAATTCCTTTATTTGCTTGATCAATATTATTAACTACGATTTTGTTACTTTTGTCATTACCAGTTTTAGCTGGAGCTATTACTATATTACTTTTTGATCGAAATTTAAA

>11750-Sycophila_biguttata-HCO

AAAGATATTGGAGTTTTATATTTAATTTTTGGGATATGAAGTGGAGTTTTAGGTTTATCTTTAAGAATATTAATTCGGTTAGAATTAGGAAATCCTGGTTCTTTAATTGGTAATGATCAAATTTATAATTCAATTGTCACTGCTCATGCTTTTATTATAATTTTTTTTTTTGTTATACCTGTTATGATAGGGGGATTTGGGAATTATTTAATTCCTTTAATTTTAGGAATTCCTGATATAGCTTTTCCTCGAATAAATAATATAAGATTTTGATTATTAATTCCTAGATTATTTTTATTACTTTCAAGTATATTTGTAGGTTCGGGAACTGGAACTGGATGGACCGTTTATCCCCCTTTATCTGGGAATTTATCTCATGGTGGACCATCAGTTGATTTATCTATTTTTTCTCTTCATTTAGCTGGTATTAGTTCAATTATAGGTTCAGTTAATTTTATTTCTACAATTTTGAATATAAAAATTTTTAAAATTGAGTTAATTCCTTTATTTGCTTGATCAATATTATTAACTACGATTTTGTTACTTTTGTCATTACCAGTTTTAGCCGGAGCTATTACTATATTACTTTTTGATCGAAATTTAAA

>17144-Torymus_flavipes-HCO_PUC

GATATTGGTATTTTATATTTTATTTTTGGAATATGAGCAAGGATTATAGGTTTATCGATAAGAATAATTATTCGTTTGGAATTAGGGAATCCCGGTTCCTTAATTGGTAAAGATCAGAATTATAATTTTATTGTTACTACTCATGCTTTTACTATAATTTTTTTTTTTGTTATACCTGTAATAATAGGAGGATTTGGTAATTATTTGGTACCTTTATTTTTAGGAACTCCTGATATAGCATTCCCTCGAATGAATAATATAAGATTTTGATTATTACCCCCTAGATTAATTTTATTAATTTCTAGTATATTTGTGGGTAGGGGTACTGGTACAGGGTGAACAGTTTATCCTCCTCTTTCAGGTAATTTATCTCATGGGGGTCCATCTGTAGATTTATCAATTTTTTCATTACATGTAGCTGGTTTATCATCTATTATAGGATCAATTAATTTTATTACTACAATTTTAAATATAAAGTTATTTAATCTTGAAATTATTCCTTTATTTAGTTGGGCTATATTATTAACTGCTATTTTATTATTATTATCTTTACCAGTTTTAGCTGGTGCTATTACTATATTGTTATTTGATCGTAATCTAAATACTTCA

>17145-Torymus_flavipes-HCO_PUC

GATATTGGTATTTTATATTTTATTTTTGGAATATGAGCAAGGATTATAGGTTTATCGATAAGAATAATTATTCGTTTGGAATTAGGGAATCCCGGTTCCTTAATTGGTAAAGATCAGAATTATAATTTTATTGGTCATACTCATGCTTTTACTATAATTTTTTTTTTTGTTATACCTGTAATAATAGGAGGATTTGGTAATTATTTGGTACCTTTATTTTTAGGAACTCCTGATATAGCATTCCCTCGAATGAATAATATAAGATTTTGATTATTACCCCCTAGATTAATTTTATTAATTTCTAGTATATTTGTGGGTAGGGGTACTGGTACAGGGTGAACAGTTTATCCTCCTCTTTCAGGTAATTTATCTCATGGGGGTCCATCTGTAGATTTATCAATTTTTTCATTACATGTAGCTGGTTTATCATCTATTATAGGATCAATTAATTTTATTACTACAATTTTAAATATAAAGTTATTTAATCTTGAAATTATTCCTTTATTTAGTTGGGCTATATTATTAACTGCTATTTTATTATTATTATCTTTACCAGTTTTAGCTGGTGCTATTACTATATTGTTATTTGATCGTAATCTAAATACTTCA

>17147-Torymus_flavipes-HCO_PUC

GATATTGGTATTTTATATTTTATTTTTGGAATATGAGCAAGGATTATAGGTTTATCGATAAGAATAATTATTCGTTTGGAATTAGGGAATCCCGGTTCCTTAATTGGTAAAGATCAGAATTATAATTTTATTGGTCCTACTCATGCTTTTACTATAATTTTTTTTTTTGTTATACCTGTAATAATAGGAGGATTTGGTAATTATTTGGTACCTTTATTTTTAGGAACTCCTGATATAGCATTCCCTCGAATGAATAATATAAGATTTTGATTATTACCCCCTAGATTAATTTTATTAATTTCTAGTATATTTGTGGGTAGGGGTACTGGTACAGGGTGAACAGTTTATCCTCCTCTTTCAGGTAATTTATCTCATGGGGGTCCATCTGTAGATTTATCAATTTTTTCATTACATGTAGCTGGTTTATCATCTATTATAGGATCAATTAATTTTATTACTACAATTTTAAATATAAAGTTATTTAATCTTGAAATTATTCCTTTATTTAGTTGGGCTATATTATTAACTGCTATTTTATTATTATTATCTTTACCAGTTTTAGCTGGTGCTATTACTATATTGTTATTTGATCGTAATCTAAATACTTCA

>17151-Torymus_sp-HCO_PUC

GATATTGGTATTTTATATTTTATTTTTGGAATATGGGCAGGAATTATAGGATTATCAATAAGAATAATTATTCGTTTAGAATTAGGAACTCCTGGTTCCTTAATTGGTAATGATCAAATTTATAATTCTATTGTTACTACTCATGCTTTTACTATAATTTTTTTTTTTGTTATACCTGTTATAATAGGAGGTTTTGGTAATTATTTAATTCCTTTATTTTTAGGAGCCCCTGATATGGCCTTTCCTCGAATAAATAATATAAGATTTTGATTATTACCTCCTAGCTTAATTTTATTAATTTCTAGAATATTTGTAGGTAGTGGTACTGGAACAGGTTGAACTGTTTATCCTCCTCTTTCAGGTAATTTATCTCATGGGGGTCCATCAGTAGATTTATCAATTTTTTCTTTGCATATTGCTGGTTTATCCTCAATTATGGGATCAATTAATTTTATTTCAACTATTATTAATATAAAATTATTTAATATTGAGATTATTCCTTTATTTAGGTGAGCCATATTATTAACAGCTATTTTATTATTATTATCTTTACCTGTGTTAGCAGGGGCTATTACTATATTATTATTTGATCGTAATTTAAATACTTCT

>17154-Torymus_auratus-HCO_PUC

GATATTGGTATTTTATATTTTATTTTTGGAATATGAGCAGGAATTATAGGATTATCTATGAGTATAATTATTCGTTTGGAGTTAGGAACTCCTGGGTCTTTAATTGGGAATGATCAAATTTATAATTCTATTGTTACTACTCATGCTTTTACTATAATTTTTTTTTTTGTTATACCTGTTATAATAGGTGGTTTCGGTAATTATTTGATTCCTTTATTTTTAGGAGTTCCTGATATAGCTTTTCCTCGAATAAATAATATGAGTTTTTGATTATTACCTCCTAGAATTATTTTATTAATTTCTAGAATATTTGTAGGGTCAGGAACGGGAACTGGTTGAACTGTTTATCCTCCTTTATCTGGCAATTTATCTCATGGGGGTCCGTCAGTTGATTTATCAATTTTTTCTTTACATGTTGCTGGACTTTCATCTATTATAGGATCAATTAATTTTATTACTACTATTTTAAATATAAAGTTATTTACTCTTGAAATTATTCCTTTATTTAGATGAGCTATATTATTAACTGCAATTTTATTACTATTGTCCTTACCAGTATTAGCTGGTGCTATTACTATATTATTATTTGATCGTAATTTAAATACTTCA

>17156-Torymus_auratus-HCO_PUC

GATATTGGTATTTTATATTTTATTTTTGGAATATGAGCAGGAATTATAGGATTATCTATGAGAATAATTATTCGTTTGGAGTTAGGAACTCCTGGGTCTTTAATTGGGAATGATCAAATTTATAATTCTATTGTTACTACTCATGCTTTTACTATAATTTTTTTTTTTGTTATACCTGTTATAATAGGTGGTTTCGGTAATTATTTGATTCCTTTATTTTTAGGAGTCCCTGATATAGCTTTTCCTCGAATAAATAATATGAGTTTTTGATTATTACCCCCTAGAATTATTTTATTAATTTCTAGAATATTTGTAGGGTCGGGAACGGGAACTGGTTGGACTGTTTATCCTCCTTTATCTGGCAATTTATCTCATGGGGGTCCGTCAGTTGATTTATCAATTTTTTCTTTACATGTTGCTGGACTTTCATCTATTATAGGATCAATTAATTTTATTACTACTATTTTAAATATAAAGTTATTTACTCTTGAAATTATTCCTTTATTTAGATGAGCTATATTATTAACTGCAATTTTATTATTATTGTCTTTACCAGTATTAGCTGGTGCTATTACTATATTATTATTTGATCGTAATTTAAATACTTCA

>17157-Torymus_rubi-HCO_PUC

GATATTGGTATTTTATATTTTATTTTTGGTATATGAGCAGGCATTATGGGATTATCGATAAGAATAATTATTCGTTTAGAACTTGGTACTCCTGGGTCTTTAATTGGTAATGATCAAATTTATAATTCTATTGTCACTACTCATGCTTTTACTATAATTTTTTTTTTTGTTATACCTGTTATGATAGGAGGTTTTGGTAATTATTTAATTCCTTTATTTTTAGGTGTTCCTGATATAGCTTTTCCTCGTATAAATAATATAAGATTTTGATTATTACCTCCTAGAATTATTTTATTAATTTCTAGAATATTTGTTGGTTCAGGTACTGGTACTGGTTGAACAGTATATCCTCCATTATCAGGTAATCTTTCTCATGGTGGTCCATCAGTTGATTTATCTATTTTTTCTTTACATGTTGCTGGTTTATCATCAATTATAGGTTCTATTAATTTTATTACTACTATTTTAAATATAAAATTATTTAATATTGAAATTATCCCTTTATTTAGGTGAGCTATATTATTAACTGCTATTTTATTATTATTATCTTTACCTGTATTAGCTGGAGCTATTACTATATTATTATTTGATCGTAATTTAAACACTTCT

>17158-Torymus_flavipes-HCO_PUC

GATATTGGTATTTTATATTTTATTTTTGGAATATGAGCAGGGATTATAGGTTTATCGATAAGAATAATTATTCGTTTGGAATTCGGGACTCCAGGTTCTTTAATTGGTAATGATCAGAATTATAATTTTATTGTTCATACTCATGCTTTTACTATAATTTTTTTTTTTGTTATACCTGTAATAATAGGAGGATTTGGTAATTATTTGGTACCTTTATTTTTAGGAACTCCTGATATAGCATTCCCTCGAATGAATAATATAAGATTTTGATTATTACCCCCTAGATTAATTTTATTAATTTCTAGTATATTTGTGGGTAGGGGTACTGGTACAGGGTGAACAGTTTATCCTCCTCTTTCAGGTAATTTATCTCATGGGGGTCCATCTGTAGATTTATCAATTTTTTCATTACATGTAGCTGGTTTATCATCTATTATAGGATCAATTAATTTTATTACTACAATTTTAAATATAAAATTATTTACTCTTGAAATTATTCCTTTATTTAGTTGGGCTATATTATTAACTGCTATTTTATTATTATTATCTTTACCAGTTTTAGCTGGTGCTATTACTATATTGTTATTTGATCGTAATCTAAATACTTCA

>17159-Torymus_flavipes-HCO_PUC

GATATTGGTATTTTATATTTTATTTTTGGAATATGAGCAGGGATTATAGGTTTATCGATAAGAATAATTATTCGTTTGGAATTAGGGACTCCTGGTTCTTTAATTGGTAATGATCAAATTTATAATTTTATTGTTACTACTCATGCTTTTACTATAATTTTTTTTTTTGTTATACCTGTAATAATAGGAGGATTTGGTAATTATTTGGTACCTTTATTTTTAGGAACTCCTGATATAGCATTCCCTCGAATGAATAATATAAGATTTTGATTATTACCCCCTAGATTAATTTTATTAATTTCTAGTATATTTGTGGGTAGGGGTACTGGTACAGGGTGAACAGTTTATCCTCCTCTTTCAGGTAATTTATCTCATGGGGGTCCATCTGTAGATTTATCAATTTTTTCATTACATGTAGCTGGTTTATCATCTATTATAGGATCAATTAATTTTATTACTACAATTTTAAATATAAAGTTATTTAATCTTGAAATTATTCCTTTATTTAGTTGGGCTATATTATTAACTGCTATTTTATTATTATTATCTTTACCAGTTTTAGCTGGTGCTATTACTATATTGTTATTTGATCGTAATCTAAATACTTCA

>17165-Torymus_geranii-HCO_PUC

GATATTGGTATTTTATATTTTATTTTTGGCATATGAGCAGGAATTATAGGTCTTTCTATAAGAATAATTATTCGTTTAGAATTGGGTACTCCAGGATCTTTAATTGGAAATGATCAAATTTATAATTCAATTGTTACTACTCATGCTTTTACTATGATTTTTTTTTTTGTTATACCTGTAATAATAGGTGGTTTTGGAAATTATTTAATTCCTTTATTTTTAGGAGTCCCAGATATAGCTTTTCCTCGGATAAATAATATAAGTTTTTGATTATTACCTCCTAGAATTATTTTATTAATTTCGAGAATATTTGTAGGATCAGGTACTGGTACAGGATGAACTGTTTATCCTCCTTTATCTGGAAATTTATCTCATGGTGGACCTTCAGTTGATTTATCAATTTTTTCTTTACATGTGGCTGGATTATCATCAATTATAGGATCGATTAATTTTATTACTACTATTTTAAATATAAAATTATTTTCTCTTGAGATTATTCCTTTATTTAGTTGAGCTATATTATTGACTGCAATTTTATTATTATTATCATTGCCTGTTTTAGCTGGGGCTATTACTATGTTATTATTTGATCGTAATTTAAATACTTCT

>17167-Torymus_auratus-HCO_PUC

GATATTGGTATTTTATATTTTATTTTTGGAATATGAGCAGGAATTATAGGATTATCTATGAGAATAATTATTCGTTTGGAGTTAGGAACTCCTGGGTCTTTAATTGGGAATGATCAAATTTATAATTCTATTGTTACTACTCATGCTTTTACTATAATTTTTTTTTTTGTTATACCTGTTATAATAGGTGGTTTCGGTAATTATTTGATTCCTTTATTTTTAGGGGTCCCTGATATAGCTTTTCCTCGAATAAATAATATGAGTTTTTGATTATTACCTCCTAGAATTATTTTATTAATTTCTAGAATATTTGTAGGGTCAGGAACGGGAACTGGTTGAACTGTTTATCCTCCTTTATCTGGCAATTTATCTCATGGGGGTCCGTCAGTTGATTTATCAATTTTTTCTTTACATGTTGCTGGACTTTCATCTATTATAGGATCAATTAATTTTATTACTACTATTTTAAATATAAAGTTATTTACTCTTGAAATTATTCCTTTATTTAGATGAGCTATATTATTAACTGCAATTTTATTATTATTATCTTTACCAGTATTAGCTGGTGCTATTACTATATTATTATTTGATCGTAATTTAAATACTTCA

>17171-Torymus_auratus-HCO_PUC

GATATTGGTATTTTATATTTTATTTTTGGAATATGAGCAGGAATTATAGGATTATCTATGAGAATAATTATTCGTTTGGAGTTAGGAACTCCTGGGTCTTTAATTGGGAATGATCAAATTTATAATTCTATTGTTACTACTCATGCTTTTACTATAATTTTTTTTTTTGTTATACCTGTTATAATAGGTGGTTTCGGTAATTATCTGATTCCTTTATTTTTAGGGGTCCCTGATATAGCTTTTCCTCGAATAAATAATATGAGTTTTTGATTATTACCTCCTAGAATTATTTTATTAATTTCTAGAATATTTGTAGGGTCAGGAACGGGAACTGGTTGAACTGTTTATCCTCCTCTATCTGGCAATTTATCTCATGGGGGTCCGTCAGTTGATTTATCAATTTTTTCTTTACATGTTGCTGGACTTTCATCTATTATAGGATCAATTAATTTTATTACTACTATTTTAAATATAAAGTTATTTACTCTTGAAATTATTCCTTTATTTAGATGAGCTATATTATTAACTGCAATTTTATTATTATTATCTTTACCAGTATTAGCTGGTGCTATTACTATATTATTATTTGATCGTAATTTAAATACTTCA

>17182-Torymus_affinis-HCO_PUC

GATATTGGTATTTTATATTTTATTTTTGGTATATGGGCTGGGATTATAGGTTTGTCTATAAGAATAATTATTCGATTGGAGTTAGGGACTCCCGGATCTTTAATTGGTAATGATCAAATTTATAATTCTATTGTAACTACTCATGCTTTTACTATAATTTTTTTTTTTGTTATACCTGTTATGATAGGGGGATTTGGGAATTATTTAATTCCTCTATTTTTAGGTGTTCCGGATATGGCTTTTCCTCGTATAAATAATATAAGGTTTTGGTTACTTCCTCCTAGAATTATTCTTTTAATTTCTAGTATATTTATTGGAACTGGTACTGGTACAGGGTGGACTGTTTATCCTCCTTTATCTGGAAATTTATCTCATGGTGGACCATCAGTTGATTTATCAATTTTTTCTTTGCATATTGCTGGTCTTTCTTCTATTATAGGGTCTATTAATTTTATTACAACAATTTTAAATATAAAGTTGTTTAATTTAGAGGTTATTCCTTTATTTAGTTGGGCCATATTATTAACTGCTATCTTATTATTATTATCTTTGCCTGTTTTAGCAGGGGCTATTACTATATTATTATTTGATCGTAATTTAAATACTTCA

>17184-Torymus_affinis-HCO_PUC

GATATTGGTATTTTATATTTTATTTTTGGTATATGGGCTGGGATTATAGGTTTGTCTATAAGAATAATTATTCGGTTGGAGTTAGGGACCCCCGGATCTTTAATTGGTAATGATCAAATTTATAATTCTATTGTAACTACTCATGCTTTTACTATAATTTTTTTTTTTGTTATACCTGTTATGATAGGGGGATTTGGGAATTATTTAATTCCTTTATTTTTAGGTGTTCCGGATATGGCTTTTCCTCGTATAAATAATATAAGGTTTTGGCTACTTCCCCCTAGAATTATTCTTTTAATTTCTAGTATATTTATTGGAACTGGTACTGGTACAGGGTGGACTGTTTATCCTCCTTTATCTGGAAATTTATCTCATGGTGGACCATCAGTTGATTTATCAATTTTTTCTTTACATATTGCTGGTCTTTCTTCTATTATAGGGTCTATTAATTTTATTACAACAATTTTAAATATAAAATTGTTTAATTTAGAGGTTATTCCTTTATTTAGTTGGGCCATATTATTAACTGCTATCTTATTATTATTATCTTTACCTGTTTTAGCAGGGGCTATTACTATATTATTATTTGATCGTAATTTAAATACTTCA

>17185-Torymus_affinis-HCO_PUC

GATATTGGTATTTTATATTTTATTTTTGGTATATGGGCTGGGATTATAGGTTTGTCTATAAGAATAATTATTCGGTTGGAGTTAGGGACCCCCGGATCTTTAATTGGTAATGATCAAATTTATAATTCTATTGTAACTACTCATGCTTTTACTATAATTTTTTTTTTTGTTATACCTGTTATGATAGGGGGATTTGGGAATTATTTAATTCCTCTATTTTTAGGTGTTCCGGATATGGCTTTTCCTCGTATAAATAATATAAGGTTTTGGTTACTTCCTCCTAGAATTATTCTTTTAATTTCTAGTATATTTATTGGAACTGGTACTGGTACAGGGTGGACTGTTTATCCCCCTTTATCTGGAAATTTATCTCATGGTGGACCATCAGTTGATTTATCAATTTTTTCTTTACATATTGCTGGTCTTTCTTCTATTATAGGGTCTATTAATTTTATTACAACAATTTTAAATATAAAGTTGTTTAATTTAGAGGTTATTCCTTTATTTAGTTGGGCCATATTATTAACTGCTATCTTATTATTATTATCTTTACCTGTTTTAGCAGGGGCTATTACTATATTATTATTTGATCGTAATTTAAATACTTCA

>17186-Torymus_affinis-HCO_PUC

GATATTGGTATTTTATATTTTATTTTTGGTATATGAGCCGGAATTATAGGTTTATCTATAAGAATAATTATTCGGTTAGAGTTAGGGACTCCAGGGTCCTTAATTGGGAATGATCAGATCTATAATTTTATTGTAACTACTCATGCTTTTACTATAATTTTTTTTTTTGTTATACCTGTTATGATAGGGGGGTTTGGGAATTATTTAATTCCTTTATTTTTAGGTGTTCCGGATATGGCTTTTCCCCGTATAAATAATATAAGGTTTTGATTACTTCCTCCTAGTATTTTTCTTTTAGTTTCTAGTATATTTATTGGAACCGGTACTGGTACAGGGTGAACTGTTTACCCTCCTTTATCTGGAAATTTATCTCATGGTGGACCATCAGTTGATTTATCAATTTTTTCTTTGCATATTGCTGGGCTTTCTTCTATTATAGGGTCTATTAATTTTATTACAACAATTTTAAATATAAAATTGTTTAATTTAGAGGTTATTCCTTTATTTAGTTGGGCTATATTATTAACTGCTATTTTATTATTATTATCTTTACCTGTTTTAGCAGGGGCTATTACTATGTTATTATTTGATCGTAATTTAAATACTTCA

>17188-Torymus_flavipes-HCO_PUC

GATATTGGTATTTTATATTTTATTTTTGGAATATGAGCAAGGATTATAGGTTTATCGATAAGAATAATTATTCGTTTGGAATTAGGGAATCCCGGTTCCTTAATTGGTAAAGATCAGAATTATAATTTTATTGGTCCTACTCATGCTTTTACTATAATTTTTTTTTTTGTTATACCTGTAATAATAGGAGGATTTGGTAATTATTTGGTACCTTTATTTTTAGGAACTCCTGATATAGCATTCCCTCGAATGAATAATATAAGATTTTGATTATTACCCCCTAGATTAATTTTATTAATTTCTAGTATATTTGTGGGTAGGGGTACTGGTACAGGGTGAACAGTTTATCCTCCTCTTTCAGGTAATTTATCTCATGGGGGTCCATCTGTAGATTTATCAATTTTTTCATTACATGTAGCTGGTTTATCATCTATTATAGGATCAATTAATTTTATTACTACAATTTTAAATATAAAGTTATTTAATCTTGAAATTATTCCTTTATTTAGTTGGGCTATATTATTAACTGCTATTTTATTATTATTATCTTTACCAGTTTTAGCTGGTGCTATTACTATATTGTTATTTGATCGTAATCTAAATACTTCA

>17190-Torymus_geranii-HCO_PUC

GATATTGGTATTTTATATTTTATTTTTGGCATATGAGCAGGAATTATAGGTCTTTCTATAAGAATAATTATTTGTTTAGAATTGGGTACTCCCGGATCTTTAATTGGAAATGATCAAATTTATAATTCAATTGTTACTACTCATGCTTTTACTATGATTTTTTTTTTTGTTATACCTGTAATAATAGGTGGTTTTGGAAATTATTTAATTCCTTTATTTTTAGGAGTCCCAGATATAGCTTTTCCTCGGATAAATAATATAAGTTTTTGATTATTACCTCCTAGAATTATTTTATTAATTTCGAGAATATTTGTAGGATCAGGTACTGGTACAGGATGAACTGTTTATCCTCCTTTATCTGGAAATTTATCTCATGGTGGACCTTCAGTTGATTTATCAATTTTTTCTTTACATGTGGCTGGATTATCATCAATTATAGGATCGATTAATTTTATTACTACTATTTTAAATATAAAATTATTTTCTCTTGAGATTATTCCTTTATTTAGTTGAGCTATATTATTGACTGCAATTTTATTATTATTATCATTGCCTGTTTTAGCTGGGGCTATTACTATGTTATTATTTGATCGTAATTTAAATACTTCT

>17192-Torymus_bedeguaris-HCO_PUC

GATATTGGTATTTTATATTTTATTTTTGGTATATGATCTGGTATTATAGGATTATCAATAAGAATGATTATTCGTTTAGAATTAGGTACCCCTGGGTCATTAATTGGTAATGATCAGATTTATAATTCTATTGTTACTATTCATGCTTTTACTATAATTTTTTTTTTTGTTATACCTGTAATAATAGGTGGTTTTGGAAATTTTTTTATTCCTTTATTTTTAGGGGTTCCTGATATAGCTTTTCCTCGAATGAATAATATAAGTTTTTGATTATTACCCCCTAGATTAATTTTATTAATTTCCAGGATATTCATTGGAAGGGGTTCTGGAACTGGTTGAACTGTTTATCCTCCATTATCTGGCAATTTATCTCATAGAGGTCCTTCAGTTGATTTATCTATTTTTTCATTACATATTGCAGGTATGTCTTCTATTATGGGATCAATTAATTTTATTTCAACTATTATAAATATAAAATTGTTTACTTTTGAGATTATTCCTTTGTTTAGTTGAGCTATATTATTAACTGCTATTTTATTATTATTATCATTACCTGTATTAGCAGGTGCAATTACTATATTGTTATTTGATCGTAATTTAAATACTTCT

>17196-Torymus_bedeguaris-HCO_PUC

GATATTGGTATTTTATATTTTATTTTTGGTATATGATCTGGTATTATAGGACTATCAATAAGAATGATTATTCGTTTAGAATTAGGTACCCCTGGGTCATTAATTGGTAATGATCAGATTTATAATTCTATTGTTACTATTCATGCTTTTACTATAATTTTTTTTTTTGTTATACCTGTAATAATAGGTGGTTTTGGAAATTTTTTTATTCCTTTATTTTTAGGGGTTCCTGATATAGCTTTTCCTCGAATGAATAATATAAGTTTTTGATTATTACCCCCTAGATTAATTTTATTAATTTCTAGGATATTTATTGGAAGGGGTTCTGGAACTGGTTGAACTGTTTATCCTCCATTATCTGGCAATTTATCTCATAGAGGTCCTTCAGTTGATTTATCTATTTTTTCATTACATATTGCAGGTATGTCTTCTATTATGGGATCAATTAATTTTATTTCAACTATTATAAATATAAAATTGTTTACTTTTGAGATTATATCTTTGTTTAGTTGAGCTATATTATTAACTGCTATTTTATTATTATTATCATTACCTGTATTAGCAGGTGCAATTACTATATTGTTATTTGATCGTAATTTAAATACTTCT

>17198-Torymus_bedeguaris-HCO_PUC

GATATTGGTATTTTATATTTTATTTTTGGTATATGATCTGGTATTATAGGATTATCAATAAGAATGATTATTCGTTTAGAATTAGGTATCCCTGGGTCATTAATTGGTAATGATCAGATTTATAATTCTATTGTTACTATTCATGCTTTTACTATAATTTTTTTTTTTGTTATACCTGTAATAATAGGTGGTTTTGGAAATTTTTTTATTCCTTTATTTTTAGGGGTTCCTGATATAGCTTTTCCTCGAATGAATAATATAAGTTTTTGATTATTACCCCCTAGATTAATTTTATTAATTTCTAGGATATTTATTGGAAGGGGTTCTGGAACTGGTTGAACTGTTTATCCTCCATTATCTGGCAATTTATCTCATAGAGGCCCTTCAGTTGATTTATCTATTTTTTCATTACATATTGCAGGTATGTCTTCTATTATGGGATCAATTAATTTTATTTCAACTATTATAAATATAAAATTGTTTACTTTTGAGATTATTCCTTTGTTTAGTTGAGCTATATTATTAACTGCTATTTTATTATTATTATCATTACCTGTATTAGCAGGTGCAATTACTATATTGTTATTTGATCGTAATTTAAATACTTCT

>17205-Torymus_rubi-HCO_PUC

GATATTGGTATTTTATATTTTATTTTTGGTATATGAGCAGGTATTATGGGATTATCGATAAGAATAATTATTCGTTTAGAACTTGGTACTCCTGGGTCTTTAATTGGTAATGATCAAATTTATAATTCTATTGTTACTACTCATGCTTTTACTATAATTTTTTTTTTTGTCATACCTGTTATGATAGGAGGTTTTGGTAATTATTTAATTCCTTTATTTTTAGGTGTTCCTGATATAGCTTTTCCTCGTATAAACAATATAAGATTTTGATTATTACCTCCTAGAATTATTTTATTAATTTCTAGAATATTTGTTGGTTCAGGTACTGGTACTGGTTGAACAGTATATCCTCCATTATCAGGTAATCTTTCTCATGGTGGTCCATCAGTTGACTTATCTATTTTTTCTTTACATGTTGCTGGTTTATCATCAATTATAGGTTCTATTAATTTTATTACTACTATTTTAAATATAAAATTATTTAATATTGAAATTATTCCTTTATTTAGTTGAGCTATATTATTAACTGCTATTTTATTATTATTATCTTTACCTGTATTAGCTGGAGCTATTACTATATTATTATTTGATCGTAATTTAAACACTTCT

>17206-Torymus_rubi-HCO_PUC

GATATTGGTATTTTATATTTTATTTTTGGTATATGAGCAGGTATTATGGGATTATCGATAAGAATAATTATTCGTTTAGAACTTGGTACTCCTGGGTCTTTAATTGGTAATGATCAAATTTATAATTCTATTGTCACTACTCATGCTTTTACTATAATTTTTTTTTTTGTTATACCTGTTATGATAGGAGGTTTTGGTAATTATTTAATTCCTTTATTTTTAGGTGTTCCTGATATAGCTTTTCCTCGTATAAATAATATAAGATTTTGATTATTACCTCCTAGAATTATTTTATTAATTTCTAGAATATTTGTTGGTTCAGGTACTGGTACTGGTTGAACAGTATATCCTCCATTATCAGGTAATCTTTCTCATGGTGGTCCATCAGTTGATTTATCTATTTTTTCTTTACATGTTGCTGGTTTATCATCAATTATAGGTTCTATTAATTTTATTACTACTATTTTAAATATAAAATTATTTAATATTGAAATTATTCCTTTATTTAGTTGAGCTATATTATTAACTGCTATTTTATTATTATTATCTTTACCTGTATTAGCTGGAGCTATTACTATATTATTATTTGATCGTAATTTAAACACTTCT

>17923-Sycophila_biguttata-HCO

AAAGATATTGGAGTTTTATATTTAATTTTTGGAATATGAAGTGGAGTTTTAGGTTTATCTTTAAGAATATTAATTCGGTTAGAATTAGGAAATCCTGGTTCTTTAATTGGTAATGATCAAATTTATAATTCAATTGTCACTGCTCATGCTTTTATTATAATTTTTTTTTTTGTTATACCTGTTATGATAGGGGGATTTGGAAATTATTTAATTCCTTTAATTTTAGGAATTCCTGATATAGCTTTTCCTCGAATAAATAATATAAGATTTTGATTATTAATTCCTAGATTATTTTTATTACTTTCAAGTATATTTGTAGGTTCGGGTACTGGGACTGGGTGGACCGTTTATCCACCTTTATCTGGGAATTTATCTCATGGTGGACCATCAGTTGATTTATCTATTTTTTCTCTTCATTTAGCTGGTATTAGTTCAATTATAGGTTCAGTCAATTTTATTTCTACAATTTTGAATATAAAAATTTTTAAAATTGAGTTAATTCCTTTATTTGCTTGATCAATATTATTAACTACGATTTTGTTACTTTTGTCATTACCAGTTTTAGCTGGAGCTATTACTATATTACTTTTTGATCGAAATTTAAA

>17924-Sycophila_biguttata-HCO_PUC

AAAGATATTGGAGTTTTATATTTAATTTTTGGGATATGAAGTGGAGTTTTAGGTTTATCTTTAAGAATATTAATTCGGTTAGAATTAGGAAATCCTGGTTCTTTAATTGGTAATGATCAAATTTATAATTCAATTGTCCCTGCTCATGCTTTTATTATAATTTTTTTTTTTGTTATACCTGTTATGATAGGGGGATTTGGGAATTATTTAATTCCTTTAATTTTAGGAATTCCTGATATAGCTTTTCCTCGAATAAATAATATAAGATTTTGATTATTAATTCCTAGATTATTTTTATTACTTTCAAGTATATTTGTAGGTTCGGGAACTGGAACTGGATGGACCGTTTATCCACCTTTATCTGGGAATTTATCTCATGGTGGACCATCAGTTGATTTATCTATTTTTTCTCTTCATTTAGCTGGTATTAGTTCAATTATAGGTTCAGTTAATTTTATTTCTACAATTTTGAATATAAAAATTTTTAAAATTGAGTTAATTCCTTTATTTGCTTGATCAATATTATTAACTACGATTTTGTTACTTTTGTCATTACCAGTTTTAGCCGGAGCTATTACTATATTACTTTTTGATCGAAATTTAAA

>17931-Sycophila_biguttata-HCO_PUC

AAAGATATTGGAGTTTTATATTTAATTTTTGGGATATGAAGTGGAGTTTTAGGTTTATCTTTAAGAATATTAATTCGGTTAGAATTAGGAAATCCTGGTTCTTTAATTGGTAATGATCAAATTTATAATTCAATTGTCCCTGCTCATGCTTTTATTATAATTTTTTTTTTTGTTATACCTGTTATGATAGGGGGATTTGGAAATTATTTAATTCCTTTAATTTTAGGAATTCCTGATATAGCTTTTCCTCGAATAAATAATATAAGATTTTGATTATTAATTCCTAGATTATTTTTATTACTTTCAAGTATATTTGTAGGTTCGGGAACTGGAACTGGATGGACCGTTTATCCACCTTTATCTGGAAATTTATCTCATGGTGGACCATCAGTTGATTTATCTATTTTTTCTCTTCATTTAGCTGGAATTAGTTCAATTATAGGTTCAGTTAATTTTATTTCTACAATTTTGAATATAAAAATTTTTAAAATTGAGTTAATTCCTTTATTTGCTTGATCAATATTATTAACTACGATTTTGTTACTTTTGTCATTACCAGTTTTAGCTGGAGCTATTACTATATTACTTTTTGACCGAAATTTAAA

>17932-Sycophila_biguttata-HCO

AAAGATATTGGAGTTTTATATTTAATTTTTGGGATATGAAGTGGAGTTTTAGGTTTATCTTTAAGAATATTAATTCGGTTAGAATTAGGAAATCCTGGTTCTTTAATTGGTAATGATCAAATTTATAATTCAATTGTCACTGCTCATGCTTTTATTATAATTTTTTTTTTTGTTATACCTGTTATGATAGGGGGATTTGGAAATTATTTAATTCCTTTAATTTTAGGAATTCCTGATATAGCTTTTCCTCGAATAAATAATATAAGATTTTGATTATTAATTCCTAGATTATTTTTATTACTTTCAAGTATATTTGTAGGTTCGGGAACTGGAACTGGATGGACCGTTTATCCACCTTTATCTGGAAATTTATCTCATGGTGGACCATCAGTTGATTTATCTATTTTTTCTCTTCATTTAGCTGGTATTAGTTCAATTATAGGTTCAGTTAATTTTATTTCTACAATTTTGAATATAAAAATTTTTAAAATTGAGTTAATTCCTTTATTTGCTTGATCAATATTATTAACTACGATTTTGTTACTTTTGTCATTACCAGTTTTAGCTGGAGCTATTACTATATTACTTTTTGACCGAAATTTAAA

>17933-Sycophila_biguttata_HCO

AAAGATATTGGAGTTTTATATTTAATTTTTGGGATATGAAGTGGAGTTTTAGGTTTATCTTTAAGAATATTAATTCGGTTAGAATTAGGAAATCCTGGTTCTTTAATTGGTAATGATCAAATTTATAATTCAATTGTCACTGCTCATGCTTTTATTATAATTTTTTTTTTTGTTATACCTGTTATGATAGGGGGATTTGGAAATTATTTAATTCCTTTAATTTTAGGAATTCCTGACATAGCTTTTCCTCGAATAAATAATATAAGATTTTGATTATTAATTCCTAGATTATTTTTATTACTTTCAAGTATATTTGTAGGTTCGGGAACTGGAACTGGATGGACCGTTTATCCACCTTTATCTGGGAATTTATCTCATGGTGGGCCATCAGTTGATTTATCTATTTTTTCTCTTCATTTAGCTGGTATTAGTTCAATTATAGGTTCAGTTAATTTTATTTCTACAATTTTGAATATAAAAATTTTTAAAATTGAGTTAATTCCTTTATTTGCTTGATCAATATTATTAACTACGATTTTATTACTTTTGTCATTACCAGTTTTAGCTGGAGCTATTACTATATTACTTTTTGATCGAAATTTAAA

>28648-Torymus_rubi-HCO

GATATTGGTATTTTATATTTTATTTTTGGTATATGAGCAGGCATTATGGGATTATCGATAAGAATAATTATTTGTTTAGAACTTGGTACTCCTGGGTCTTTAATTGGTAATGATCAAATTTATAATTTTATTGTCACTACTCATGCTTTTACTATAATTTTTTTTTTTGTCATACCTGTTATGATAGGAGGGTTTGGTAATTATTTAATTCCTTTATTTTTAGGTGTCCCTGACATAGCTTTTCCTCGTATAAATAATATAAGATTTTGATTATTACCTCCTAGAATTATTTTATTAATTTCTAGAATATTTGTTGGTTCAGGTACTGGTACTGGTTGAACAGTATATCCTCCATTATCAGGTAATCTTTCTCATGGTGGTCCATCAGTTGATTTATCTATTTTTTCTTTACATGTTGCTGGTTTATCATCAATTATAGGTTCTATTAATTTTATTACTACTATTTTAAATATAAAATTATTTAATATTGAAATTATCCCTTTATTTAGTTGAGCTATATTATTAACTGCTATTTTATTATTATTATCTTTACCTGTATTAGCTGGAGCTATTACTATATTATTATTTGATCGTAATTTAAACACTTCT

>29628-Aulogymnus_arsames-HCO_PUC

TATTTTTGGGATATGGGCTGGAATTTTAGGATTATCAATAAGATTAATAATTTGTTTAGAATTAGGAAATCCCGGATCTTTAATTGGAAATGATCAGATTTATAATTCTATTGTTACTACTCATGCATTTACTATAATTTTTTTTTTTGTAATACCAGTAATAATAGGAGGGTTTGGAAATTATTTAATTCCTTTAATATTAGGGGTTCCAGATATAGCATTTCCCCGGATAAATAATATAAGATTTTGATTGCTTCCTCCTAGATTAATATTATTAATTTCTAGAATATTTATTGGCTCTGGGACAGGTACAGGATGAACTGTTTATCCTCCTTTATCTTCAAATTTAGGTCATAGAGGTCCTTCAGTTGATTTATCAATTTTTTCTTTGCATATTGCAGGGGCATCATCAATTATAGGGTCTATTAATTTTATTAGGACAATTTTAAATATAAAAAATTATAAAATAGAAAATATTTCTTTATTTTCATGATCAATATTATTAACTGCAATTTTATTATTATTATCTCTTCCTGTATTGGCTGGGGCTA

>29636-Aulogymnus_obscuripes-HCO

TATTTTTGGTATGTGAGCTGGGGTATTAGGTTTATCATTAAGGTTAATGATTCGATTGGAGTTAGGGAATCCCGGTTCTTTAATTGGGAATGATCAAATTTATAATTCTATTGTTACAACTCATGCTTTTACTATAATTTTCTTTTTTGTAATACCGGTTATAATAGGAGGGTTTGGAAATTATTTAATCCCTTTAATGTTAGGTACTCCGGATATAGCTTTTCCTCGAATAAATAATATAAGATTTTGGTTATTACCTCCAAGATTAACTTTATTAATTTCTAGAATATTTATTGGTTCAGGGACTGGTACAGGATGAACTGTATATCCCCCTTTATCATCAAATTTAGGTCATAGGGGTCCTTCAGTTGATTTATCTATTTTTTCTTTACATATTGCTGGGGCATCTTCAATTATAGGTTCAATTAATTTTATTACTACAATCTTAAATATAAAAAATTATAAAATAGAAAATATTTCTTTATTTTCATGATCAATGTTACTTACTGCTATTTTATTATTACTTTCATTACCGGTTTTAGCTGGGGCAA

>29638-Aulogymnus_obscuripes-HCO

TATTTTTGGTATGTGAGCTGGGGTATTAGGTTTATCATTAAGGTTAATGATTCGATTGGAGTTAGGGAATCCCGGTTCTTTAATTGGGAATGATCAAATTTATAATTCTATTGTTACAACTCATGCTTTTACTATAATTTTCTTTTTTGTAATACCGGTTATAATAGGAGGGTTTGGAAATTATTTAATCCCTTTAATGTTAGGTACTCCGGATATAGCTTTTCCTCGAATAAATAATATAAGATTTTGGTTATTACCTCCAAGATTAACTTTATTAATTTCTAGAATATTTATTGGTTCAGGGACTGGCACAGGATGAACTGTATATCCCCCTTTATCATCAAATTTAGGTCATAGGGGTCCTTCAGTTGATTTATCTATTTTTTCTTTACATATTGCTGGGGCGTCTTCAATTATAGGTTCAATTAATTTTATTACTACAATCTTAAATATAAAAAATTATAAAATAGAAAATATTTCTTTATTTTCATGATCAATGTTACTTACTGCTATTTTATTATTACTTTCATTACCGGTTTTAGCTGGGGCAA

>29641-Aulogymnus_obscuripes-HCO

TATTTTTGGTATGTGAGCTGGGGTATTAGGTTTATCATTAAGGTTAATGATTCGATTGGAGTTAGGGAATCCCGGTTCTTTAATTGGGAATGATCAAATTTATAATTCTATTGTTACAACTCATGCTTTTACTATAATTTTCTTTTTTGTAATACCGGTTATAATAGGAGGGTTTGGAAATTATTTAATCCCTTTAATGTTAGGTACTCCGGATATAGCTTTTCCTCGAATAAATAATATAAGATTTTGGTTATTACCTCCAAGATTAACTTTATTAATTTCTAGAATATTTATTGGTTCAGGGACTGGTACAGGATGAACTGTATATCCCCCTTTATCATCAAATTTAGGTCATAGGGGTCCTTCAGTTGATTTATCTATTTTTTCTTTACATATTGCTGGGGCATCTTCAATTATAGGTTCAATTAATTTTATTACTACAATCTTAAATATAAAAAATTATAAAATAGAAAATATTTCTTTATTTTCATGATCAATGTTACTTACTGCTATTTTATTATTACTTTCATTACCTGTTTTAGCTGGGGCAA

>29645-Aulogymnus_skianeuros-HCO

TATTTTTGGTATGTGAGCAGGGATTTTAGGATTATTTATGAGTTTAATAATTTGAATAGAATTGGGAAATCCTGGATCCTTAATTGGAAATGATCAAATTTATAATTCAATTGTTACTACTCATGCATTTACAATAATTTTTTTTTTTGTTATGCCTGTGATAATAGGGGGGTTTGGAAATTATTTAATTCCTTTAATATTAGGAACTCCTGATATAGCTTTTCCTCGAATAAATAACATAAGATTTTGATTATTACCTCCGAGATTAATTTTATTAATTTCAAGGATATTTATTGGAACTGGAACTGGAACTGGATGGACTGTTTATCCACCATTATCTTCAAATTTAGGACATAGAGGTCCATCAGTGGATTTGTCAATTTTTTCACTTCATATTGCTGGAGCTTCTTCAATTATGGGTTCAATTAATTTTATTACTACTATTTTAAATATAAAAAATTATAAGATAGAAAATGTTTCATTATTTTCTTGGTCAATATTATTAACAGCAATTCTTTTATTGCTGGCTCTTCCTGTATTAGCAGGGGCAA

>7303-Megastigmus_dorsalis-HCO

TTTATTTTTGGTATATGATCAGGTATTATTGGGTTATCTNTTAGATTAATTATTCGTATGGAATTAGGTAATCCAGGTTCTTTAATTGGAAATGATCAAATTTATAATTCAGTAGTTACTACTCATGCTTTTATTATAATTTTTTTTTTTGTTATACCAGTGATAATAGGGGGATTTGGAAATTTTTTAATTCCTTTAATTATAGGGGTTCCTGATATAGCTTTTCCTCGTATAAATAATATAAGTTTTTGACTTCTTCCCCCNAGAATTATATTATTAATTTCTAGAATATTTATTGGAAGAGGAACTGGTACTGGATGAACTGTTTATCCTCCTTTATCTTCTAATTTATCTCATAGAGGTCCGTCTGTGGATTTATCAATTTTTTCTTTACATATTGCTGGGGTTTCTTCAATTATAGCTTCAATTAATTTTATTTCTACTATTTTAAATATAAAGTTATTTAAAATTGATTTAATTCCTTTATTTTCTTGGTCTATACTATTAACTGCTATTTTATTATTATTATCTTTGCCTGTTTTAGCAGGTGCTATTACTATATTATTATTTGATCGA

>7443-Mesopolobus_sericeus-HCO

TTATTTTTGGGATATGATCCGGAGTAATAGGTTTATCTATAAGAATAATTATTCGATTAGAACTGGGTAATCCTGGGTCTTTAATTGGTAATGATCAAATTTATAATTCTATTGTTACCACTCACGCATTTACAATAATTTTTTTTTTTGTTATACCAGTAATAATAGGAGGATTTGGTAATTATTTAGTCCCAATAATTTTAGGGGCTCCAGATATAGCCTTCCCTCGAATAAATAATATAAGATTTTGATTATTACCACCTAGATTAATATTACTTATTTCTAGTATATTTATTGGATCTGGTACTGGTACTGGGTGAACTGTTTATCCCCCATTATCTTCAAATTTATCACACAGGGGGCCTTCGGTTGACCTATCAATTTTTTCTTTACATATTGCTGGAGCTTCTTCAATTATAGGATCAATTAATTTTATTACTACAGTATTAAATATAAAAATTTATAAAATTGATAATATTCCTTTATTAGCTTGATCAATATTGTTAACAGCAATTTTATTACTTCTATCTTTGCCAGTATTGGCCGGTGCTATTACTATATTACTATTTGATCGAAATCTG

>7448-Megastigmus_dorsalis-HCO

TTTATTTTTGGAATATGATCAGGTATTATTGGGTTATCTCTTAGGTTAATTATTNGTATGGAATTAGGAAATCCGGGTTCTTTAATTGGAAATGATCAAATTTATAATTCNGTAGTTACTACTCATGCCTTTATTATAATTTTTTTTTTTGTTATACCAGTGATAATAGGGGGATTCGGAAATTTTTTAATTCCTTTAATTATAGGGGTTCCTGATATAGCTTTTCCTCGTATAAATAATATAAGTTTTTGACTTCTTCCCCCAAGAATTATATTATTAATTTCTAGAATATTTATTGGAAGTGGAACTGGTACTGGATGAACTGTTTATCCTCCTTTATCTTNTAATTTATCTCATAGAGGCCCTTCTGTGGATTTATCAATTTTTTCTTTACATATTGCTGGGGTTTCTTCAATTATAGCTTCAATTAATTTTATTTCTACTATTTTAAATATAAAATTATTTAAAATTGATTTAATTCCTTTATTTTCTTGATCTATATTATTAACTGCTATTTTATTATTATTATCTTTGCCTGTTTTAGCAGGTGCTATTACTATATTATTATTTGATCGA

>7467-Megastigmus_dorsalis-HCO

TTTATTTTTGGTATATGATCAGGTATTATTGGGTTATCTCTTAGATTAATTATTTGTATGGAATTAGGTAATCCNGGTTCTTTAATTGGAAATGATCAAATTTATAATTCNGTAGTTACTACTCATGCTTTTATTATAATTTTTTTTTTTGTTATACCAGTGATAATAGGGGGATTTGGAAATTTTTTAATTCCTTTAATTATAGGGGTTCCTGATATAGCTTTTCCTCGTATAAATAATATAAGTTTTTGACTTCTTCCCCCAAGAATTATATTATTAATTTCTAGAATATTTATTGGAAGAGGAACTGGTACTGGATGAACTGTTTATCCTCCTTTATCTTCTAATTTATCTCATAGAGGTCCGTCTGTGGATTTATCAATTTTTTCTTTACATATTGCTGGGGTTTCTTCAATTATAGCTTCAATTAATTTTATTTCTACTATTTTAAATATAAAGTTATTTAAAATTGATTTAATTCCTTTATTTTCTTGGTCTATACTATTAACTGCTATTTTATTATTATTATCTTTGCCTGTTTTAGCAGGTGCTATTACTATATTATTATTTGATCGA

>7502-Megastigmus_dorsalis-HCO

TTTATTTTTGGTATATGATCAGGTATTATTGGGTTATCTCTTAGATTAATTATTNGTATGGAATTAGGTAATCCNGGTTCTTTAATTGGAAATGATCAAATTTATAATTCNGTAGTTACTACTCATGCTTTTATTATAATTTTTTTTTTTGTTATACCAGTGATAATAGGGGGATTTGGAAATTTTTTAATTCCTTTAATTATAGGGGTTCCTGATATAGCTTTTCCTCGTATAAATAATATAAGTTTTTGACTTCTTCCCCCAAGAATTATATTATTAATTTCTAGAATATTTATTGGAAGAGGAACTGGTACTGGATGAACTGTTTATCCTCCTTTATCTTCTAATTTATCTCATAGAGGTCCGTCTGTGGATTTATCAATTTTTTCTTTACATATTGCTGGGGTTTCTTCAATTATAGCTTCAATTAATTTTATTTCTACTATTTTAAATATAAAGTTATTTAAAATTGATTTAATTCCTTTATTTTCTTGGTCTATACTATTAACTGCTATTTTATTATTATTATCTTTGCCTGTTTTAGCAGGTGCTATTACTATATTATTATTTGATCGA

>7535-Megastigmus_dorsalis-HCO

TTTATTTTTGGTATATGATCAGGTATTATTGGGTTATCTCTTAGATTAATTATTCGTATGGAATTAGGTAATCCNGGTTCTTTAATTGGAAATGATCAAATTTATAATTCNGTAGTTACTACTCATGCTTTTATTATAATTTTTTTTTTTGTTATACCAGTGATAATAGGGGGATTTGGAAATTTTTTAATTCCTTTAATTATAGGGGTTCCTGATATAGCTTTTCCTCGTATAAATAATATAAGTTTTTGACTTCTTCCCCCAAGAATTATATTATTAATTTCTAGAATATTTATTGGAAGAGGAACTGGTACTGGATGAACTGTTTATCCTCCTTTATCTTCTAATTTATCTCATAGAGGTCCGTCTGTGGATTTATCAATTTTTTCTTTACATATTGCTGGGGTTTCTTCAATTATAGCTTCAATTAATTTTATTTCTACTATTTTAAATATAAAGTTATTTAAAATTGATTTAATTCCTTTATTTTCTTGGTCTATATTATTAACTGCTATTTTATTATTATTATCTTTGCCTGTTTTAGCAGGTGCTATTACTATATTATTATTTGATCGA

>7620-Torymus_auratus-HCO

GATATTGGTATTTTATATTTTATTTTTGGAATATGAGCAGGAATTATAGGATTATCTATGAGAATAATTATTCGTTTGGAGTTAGGAAATCCTGGGTCTTTAATTGGGAATGATCAAATTTATAATTCTATTGTTACTACTCATGCTTTTACTATAATTTTTTTTTTTGTTATACCTGTTATAATAGGTGGTTTCGGTAATTATTTGATTCCTTTATTTTTAGGAGTTCCTGATATAGCTTTTCCTCGAATAAATAATATGAGTTTTTGATTACTACCCCCTAGAATTATTTTATTAATTTCTAGAATATTTGTAGGATCAGGAACGGGAACTGGTTGAACTGTTTATCCTCCTTTATCTGGCAATTTATCTCATGGAGGTCCGTCAGTTGATTTATCAATTTTTTCTTTACATGTTGCTGGACTTTCATCTATTATAGGATCAATTAATTTTATTACTACTATTTTAAATATAAAGTTATTCACTCTTGAAATTATTCCTTTATTTAGATGAGCTATATTATTAACTGCAATTTTATTATTATTATCCTTACCAGTATTAGCTGGTGCTATTACTATATTATTATTTGATCGTAATTTAAATACTTCA

>7682-Torymus_auratus-HCO

GATATTGGTATTTTATATTTTATTTTTGGAATATGAGCAGGAATTATAGGATTATCTATGAGAATAATTATTCGTTTGGAGTTAGGAACTCCTGGGTCTTTAATTGGGAATGATCAAATTTATAATTCTATTGTTACTACTCATGCTTTTACTATAATTTTTTTTTTTGTTATACCTGTTATAATAGGTGGTTTCGGTAATTATTTGATTCCTTTATTTTTAGGGGTTCCTGATATAGCTTTTCCTCGAATAAATAATATAAGTTTTTGATTATTACCTCCTAGAATTATTTTATTAATTTCTAGAATATTTGTAGGGTCAGGAACGGGAACTGGTTGAACTGTTTATCCTCCTTTATCTGGCAATTTATCTCATGGGGGTCCGTCAGTTGATTTATCAATTTTTTCTTTACATGTTGCTGGACTTTCATCTATTATAGGATCAATTAATTTTATTACTACTATTTTAAATATAAAGTTATTTACTCTTGAAATTATTCCTTTATTTAGATGAGCTATATTATTAATTGCAATTTTATTATTATTATCCTTACCAGTATTAGCTGGTGCTATTACTATATTATTATTTGATCGTAATTTAAATACTTCA

>7687-Torymus_auratus-HCO

GATATTGGTATTTTATATTTTATTTTTGGAATATGAGCAGGAATTATAGGATTATCTATGAGAATAATTATTCGTTTGGAGTTAGGAACTCCTGGGTCTTTAATTGGGAATGATCAAATTTATAATTCTATTGTTACTACTCATGCTTTTACTATAATTTTTTTTTTTGTTATACCTGTTATAATAGGTGGTTTCGGTAATTATTTGATTCCTTTATTTTTAGGAGTCCCTGATATAGCTTTTCCTCGAATAAATAATATGAGTTTTTGATTATTACCTCCTAGAATTATTTTATTAATTTCTAGAATATTTGTAGGGTCAGGAACGGGAACTGGTTGAACTGTTTATCCTCCTTTATCTGGCAATTTATCTCATGGGGGTCCGTCAGTTGATTTATCAATTTTTTCTTTACATGTTGCTGGACTTTCATCTATTATAGGATCAATTAATTTTATTACTACTATTTTAAATATAAAGTTATTTACTCTTGAAATTATTCCTTTATTTAGATGAGCTATATTATTAACTGCAATTTTATTATTATTATCTTTACCAGTATTAGCTGGTGCTATTACTATATTATTATTTGATCGTAATTTAAATACTTCA

>7688-Torymus_auratus-HCO

GATATTGGTATTTTATATTTTATTTTTGGAATATGAGCAGGAATTATAGGATTATCTATGAGAATAATTATTCGTTTGGAGTTAGGAACTCCTGGGTCTTTAATTGGGAATGATCAAATTTATAATTCTATTGTTACTACTCATGCTTTTACTATAATTTTTTTTTTTGTTATACCTGTTATAATAGGTGGTTTCGGTAATTATTTGATTCCTTTATTTTTAGGAGTCCCTGATATAGCTTTCCCTCGAATAAATAATATGAGTTTTTGATTATTACCTCCTAGAATTATTTTACTAATTTCTAGAATATTTGTAGGGTCAGGAACGGGAACTGGTTGAACTGTTTATCCTCCTTTATCTGGCAATTTATCTCATGGGGGTCCGTCAGTTGATTTATCAATTTTTTCTTTACATGTTGCTGGACTTTCATCTATTATAGGATCAATTAATTTTATTACTACTATTTTAAATATAAAGTTATTTACTCTTGAAATTATTCCTTTATTTAGATGAGCTATATTATTAATTGCAATTTTATTATTATTATCCTTACCAGTATTAGCTGGTGCTATTACTATATTATTATTTGATCGTAATTTAAATACTTCA

>7807-Torymus_auratus-HCO_PUC

GATATTGGTATTTTATATTTTATTTTTGGAATATGAGCAGGAATTATAGGATTATCTATGAGAATAATTATTCGTTTGGAGTTAGGAACCCCTGGGTCTTTAATTGGGGATGATCAAATTTATAATTCTATTGTTACTACTCATGCTTTTACTATAATTTTTTTTTTTGTTATACCTGTTATAATAGGTGGTTTCGGTAATTATTTGATTCCTTTATTTTTAGGAGTCCCTGATATAGCTTTTCCTCGAATAAATAATATGAGTTTTTGATTATTACCTCCTAGAATTATTTTATTAATTTCTAGAATATTTGTAGGTTCAGGAACGGGAACTGGTTGAACTGTTTATCCTCCTTTATCTGGCAATTTATCTCATGGGGGTCCATCAGTTGATTTATCAATTTTTTCTTTACATGTTGCTGGACTTTCATCTATTATAGGATCAATTAATTTTATTACTACTATTTTAAATATAAAGTTATTTACTCTTGAAATTATTCCTTTATTTAGATGAGCTATATTATTAACTGCAATTTTATTATTATTGTCCTTACCAGTATTAGCTGGTGCTATTACTATATTATTATTTGATCGTAATTTAAATACTTCA

>7820-Torymus_flavipes-HCO_PUC

GATATTGGTATTTTATATTTTATTTTTGGAATATGAGCAGGGATTATAGGTTTATCGATAAGAATAATTATTCGTTTGGAATTAGGGACTCCTGGTTCTTTAATTGGTAATGATCAGATTTATAATTCTATTGTTACTACTCATGCTTTTACTATAATTTTTTTTTTTGTTATACCTGTAATAATAGGAGGATTTGGTAATTATTTGGTACCTTTATTTTTAGGAACTCCTGATATAGCATTCCCTCGAATGAATAATATAAGATTTTGATTATTACCTCCTAGATTAATTTTATTAATTTCTAGTATATTTGTGGGTAGGGGTACTGGTACAGGGTGAACAGTTTATCCTCCTCTTTCAGGTAATTTATCTCATGGGGGTCCATCTGTAGATTTATCAATTTTTTCATTACATGTAGCTGGTTTATCATCTATTATAGGATCAATTAATTTTATTACTACAATTTTAAATATAAAGTTATTTAATCTTGAAATTATTCCTTTATTTAGTTGGGCTATATTATTAACTGCTATTTTATTATTATTATCTTTACCAGTTTTAGCTGGTGCTATTACTATATTGTTATTTGATCGTAATCTAAATACTTCA
